# A *Gpr35* tuned gut-brain metabolic axis regulates depressive-like behavior

**DOI:** 10.1101/2023.06.11.542602

**Authors:** Lingsha Cheng, Haoqian Wu, Xiaoying Cai, Qiong Wang, Youying Zhang, Zhe Yin, Qingyuan Yan, Yuanlong Hou, Yonggui Yuan, Guangji Wang, Xueli Zhang, Haiping Hao, Xiao Zheng

## Abstract

Gene-environment interactions shape animal behavior and the susceptibility to neurobehavioral symptoms such as depression. However, little is known about the signaling pathway that integrates genetic and environmental inputs with neurobehavioral outcomes, preventing the development of targeted therapies. Here we report that *Gpr35* engages a gut microbe-to-brain metabolic pathway to modulate neuronal plasticity and depressive behavior in mice. Chronic stress decreases gut epithelial *Gpr35*, the genetic deletion of which induces despair and social impairment in a microbiome-dependent manner. We identify a dominant role for the imbalance of microbe-derived indole-3-carboxaldehyde (IAld) and indole-3-lactate (ILA) in the behavioral symptoms with *Gpr35* deficiency. Mechanistically, these bacterial metabolites counteractively modulate dendritic spine density and synaptic transmission in the nucleus accumbens. Supplementation of IAld, which is similarly decreased in depressive patients, produce anti-depressant effects in mice with stress or gut epithelial *Gpr35* deficiency. Together, these findings identify a genetics-shaped gut-brain connection underlying the susceptibility to depression and suggest a microbial metabolite-based therapeutic strategy to genetic predisposition.

## Introduction

Major depressive disorder (MDD), a heterogeneous neuropsychiatric disorder affecting more than 350 million people worldwide, is characterized by social interaction deficits, anhedonia and despair behaviors^1^. The exact causes of depressive behavior remain unclear, and are believed to be influenced by both genetic and environmental factors^2^. Although large-scale genome-wide association analyses (GWAS)-based studies have increasingly revealed disease-associated mutations^3, 4^, the biologic consequence remains incompletely understood, and limited progress has been made in causatively linking the genetic variants with behavioral outcome, especially from the perspective of gene-environment crosstalk^5^.

MDD patients are often afflicted with somatic symptoms such as gastrointestinal (GI) dysfunction. In the past decade, gut microbial dysbiosis has been extensively reported in both patients and animal models with depression, and probiotics have shown promise to alleviate the symptoms. In line with these findings, accumulating studies have shown extensive effects of gut microbiota, a common environmental factor, on neural activities and circuits involved in depressive behavior^6^. Via signaling along the gut-brain axis, an increasing list of microbial products, such as short-chain fatty acids and tryptophan catabolites, have been demonstrated to serve as signaling molecules that shape behavioral outcome^7–9^. Of note, gut microbial configuration and metabolites are subjected to the regulation by host genetic variation^10, 11^, although whether and how this genetic-microbe interaction is implicated in brain and behavior remains largely unexplored.

G-protein-coupled receptor 35 (*Gpr35*) is an orphan receptor predominantly expressed on the intestinal epithelia ^12^. Epidemiological study has linked *Gpr35* variance with the risk of inflammatory bowel disease, which is largely attributed to the modulation of gut epithelia and mucosal macrophages^13, 14^. More recently, studies from our group and others report that *Gpr35* serves as a novel regulator of gut symbiosis at this critical host-microbe interface^15, 16^, suggesting the possibility that it may regulate host pathophysiology via gut microbial signals. How *Gpr35* is involved in regulating gut-brain crosstalk and behavior at homeostasis and during stress remains elusive.

Here, we demonstrate a critical role of intestinal Gpr35 on microbial signals to the brain and depression-like behavior. *Gpr35* deficiency on the gut epithelia is sufficient to induce depression-like behavior in a microbiome-dependent manner. Interestingly, gut *Gpr35* dictates the balance of microbial metabolite analogues indole-3-carboxaldehyde (IAld) and indole-3-lactate (ILA), which exert conversing effects on neuroplasticity and depressive symptoms. We also show that IAld supplementation effectively rescues the depressive phenotypes. These findings uncover a regulatory role of *Gpr35* along the gut microbiome-brain axis and reveal new signaling metabolites in the regulation of depressive behavior, which may expand the therapeutic approach to genetic predisposition of depression.

## Results

### Gpr35 deficiency induces social deficits and depression-like behaviors in mice

The predominant expression of *Gpr35* in the gut suggests a potential role in gut-brain signaling. To understand how *Gpr35* may change under depression-relevant stressful conditions, we subjected mice to a short period of unpredictable mild stress (UMS), which was previously shown to induce depression-like behavior in mice^17^. We found that *Gpr35* transcripts were markedly decreased in the ileum and colon of stressed mice (**Fig.S1a**). No significant change, however, was observed in major brain regions (hippocampus, nucleus accumbens, medial prefrontal cortex) or the liver (**Fig.S1a**), which was indicative of a biased regulation of gut Gpr35 by stress.

To directly assess the role of *Gpr35* in behavioral control, we bred wild type (WT) and *Gpr35*-deficient (*Gpr35*^-/-^) mice (**Fig.S1b,c**), and made a comprehensive evaluation in terms of anxiety, despair and social interaction behavior in the littermates. In the open-field test (OFT), both male and female *Gpr35*^-/-^ mice showed significantly reduced entry into the central arena (**Fig.1a** and **Fig.S1d**). These mice also displayed less inclination to explore the central area, which indicated anxiogenic effects of *Gpr35* deficiency (**Fig.1a** and **Fig.S1d**). In line, *Gpr35*^-/-^ mice showed more immobility time in the tail-suspension and forced-swimming test (TST and FST, **Fig.1b** and **Fig.S1e**). Also, in the reciprocal social interaction test (SIT), *Gpr35*^-/-^ mice consistently lacked social preference for a mouse over an object (**Fig.1c** and **Fig.S1f**). Importantly, we verified that *Gpr35*^-/-^ mice did not have locomotive dysfunction (**Fig.S1g**). Furthermore, we found that, after exposure to a series of unpredictable mild stressors, *Gpr35*^-/-^ mice showed a trend of decreased central entries in the OFT and increased immobility in the FST, indicative of exacerbated anxiety and despair compared to the WT littermates (**Fig.S1h**). Collectively, these data indicate that *Gpr35* deficiency induces depression-like behavioral symptoms in both sexes of mice and predisposes to depressive behavior.

**Fig.1.**
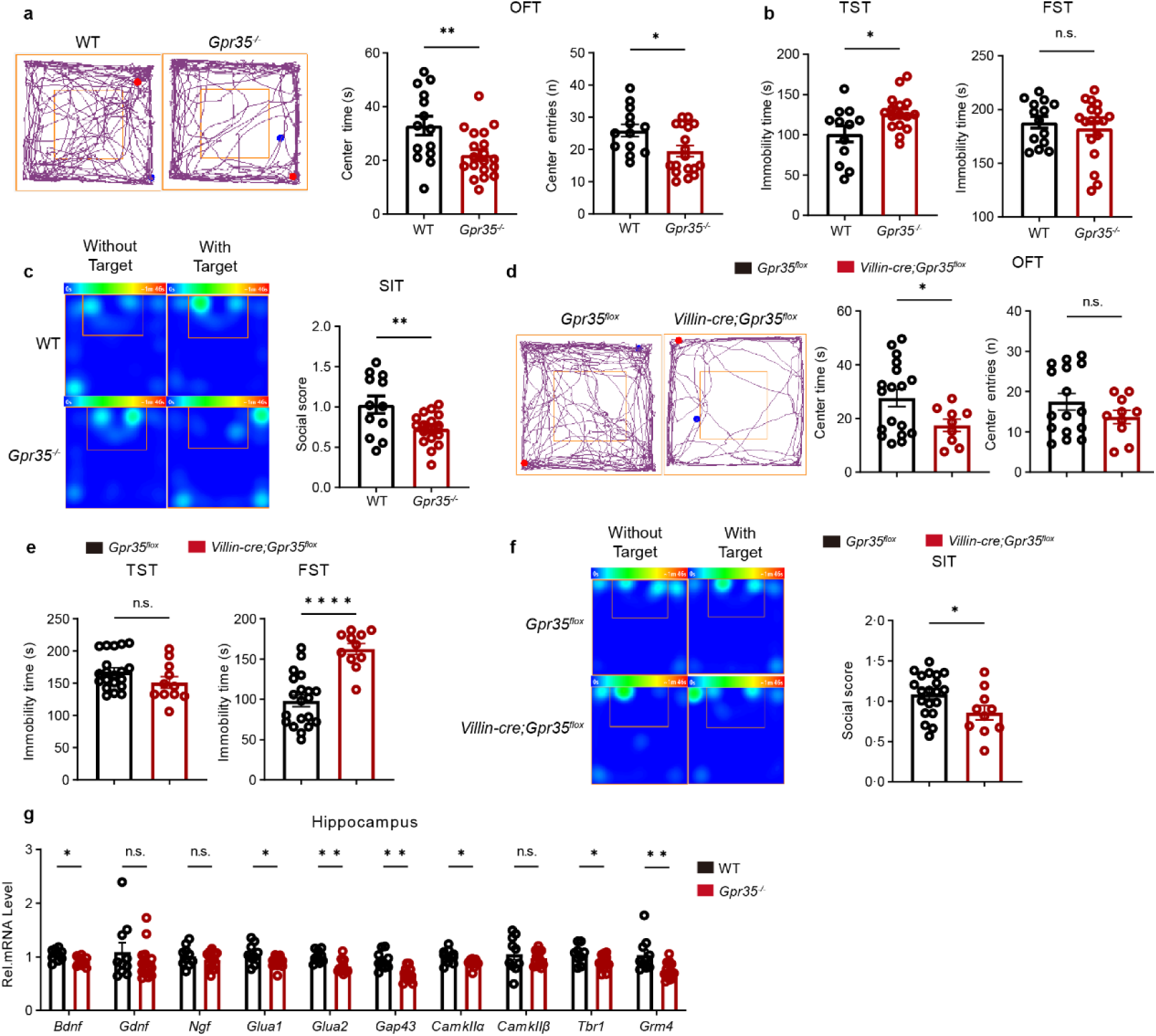
Ablation of gut epithelial *Gpr35* induces depression-like behavior in mice. (a) Anxiety-like behavior of male WT and *Gpr35*^-/-^ littermates as assessed in the open-field test (OFT). Number of entries into central area and time spent in the center were compared for n=14 and 19 mice. (b) Despair behavior of male WT and *Gpr35*^-/-^ littermates as assessed by immobility time in the tail suspension test (TST) and forced swimming test (FST) (n = 14 and 19). (c) Social interaction score of male WT and *Gpr35*^-/-^ mice as assessed in the social interaction test (SIT). (d) Anxiety-like behavior of male *Gpr35*^flox^ and *Villin-cre*;*Gpr35^flox^* littermates as assessed in the open-field test (OFT). Time spent in the center, numbers of entries and travel distance in central area were compared for n=22 and 11 mice. (e) Depression-like behavior of male *Gpr35*^flox^ and *Villin-cre*;*Gpr35^flox^* littermates as assessed by immobility time in the tail suspension test (TST) and forced swimming test (FST) (n = 16 and 10). (f) Social impairment of male *Villin*-*cre*;*Gpr35^flox^* mice as assessed in the social interaction test. (n=22 and 11). (g) Neurotrophic factor and synaptic plasticity related marker gene expression was assessed in the hippocampus isolated from WT and *Gpr35*^-/-^ mice (n=10-15). Data represent mean ± SEM. \**p* < 0.05, \*\**p* < 0.01, \*\*\*\**p* < 0.0001, n.s., no significance; two-tailed unpaired Student’s *t*-test.

To gain cell specific insights for gut epithelial *Gpr35* in behavioral modulation, we crossed *Gpr35^flox^* mice with *Villin*-*cre* mice to specifically delete *Gpr35* in gut epithelial cells (**Fig.S2a**). *Villin*-*cre*; *Gpr35^flox^* mice (male and female) showed no difference in locomotive activity (**Fig.S2b**), but displayed a spectrum of behavioral symptoms including social avoidance, despair and anxiety-like behavior, largely phenocopying those observed in *Gpr35*^-/-^ mice (**Fig.1d-f** and **Fig.S2c-f**). These results confirmed that gut *Gpr35* deficiency is sufficient to induce depression-like behavior.

We further profiled neurosynaptic and inflammatory makers in the brain to understand the cellular basis underlying the behavioral changes induced by *Gpr35* deficiency. Immunofluorescence staining of Iba-1 and GFAP revealed no significant changes in glial activation status, suggesting the absence of neuroimmune activation in *Gpr35*^-/-^ mice (**Fig.S2g,h**). Moreover, DCX^+^ staining showed comparable neurogenesis in the hippocampus of WT and *Gpr35*^-/-^ mice (**Fig.S2g,h**). Consistent with the behavioral impairment, *Gpr35*^-/-^ mice showed decreased mRNA expression of mediators involved in neuroplasticity (*Bdnf, Gap43, Tbr1*) and synaptic transmission (*Glua1, Glua2, CamKII, Grm4*) in the hippocampus (**Fig.1g**). Taken together, these findings suggest an essential role of gut Gpr35 in depression-like behavior.

### Gpr35 ablation reprograms metabolic signals from the gut microbiome

Gut barrier integrity and microbial signals play a critical role in shaping gut-brain crosstalk and behavior^18^. To understand how gut *Gpr35* deficiency induce distal neural and behavioral changes, we examined gut local changes for the primary clue. In line with our previous report^16^, we observed a significant increase in ileum and colon villi length of *Gpr35^-/-^* mice compared with the WT littermates (**Fig.S3a**). We also found reduced goblet cells density in the jejunum and ileum of *Gpr35*^-/-^ mice (**Fig.S3b**). In addition, the transcripts of cytokines and tight junction proteins showed significant changes in the colon of *Gpr35*^-/-^ mice to varying degrees (**Fig.S3c,d**). In line with the molecular changes, oral gavage of FITC-dextran confirmed impairment in gut barrier integrity of *Gpr35*^-/-^ mice (**Fig.2a**). Together, these findings suggest that *Gpr35* deficiency disturbs gut epithelial homeostasis and barrier function, which may therefore translate to altered gut-to-brain signaling.

**Fig.2.**
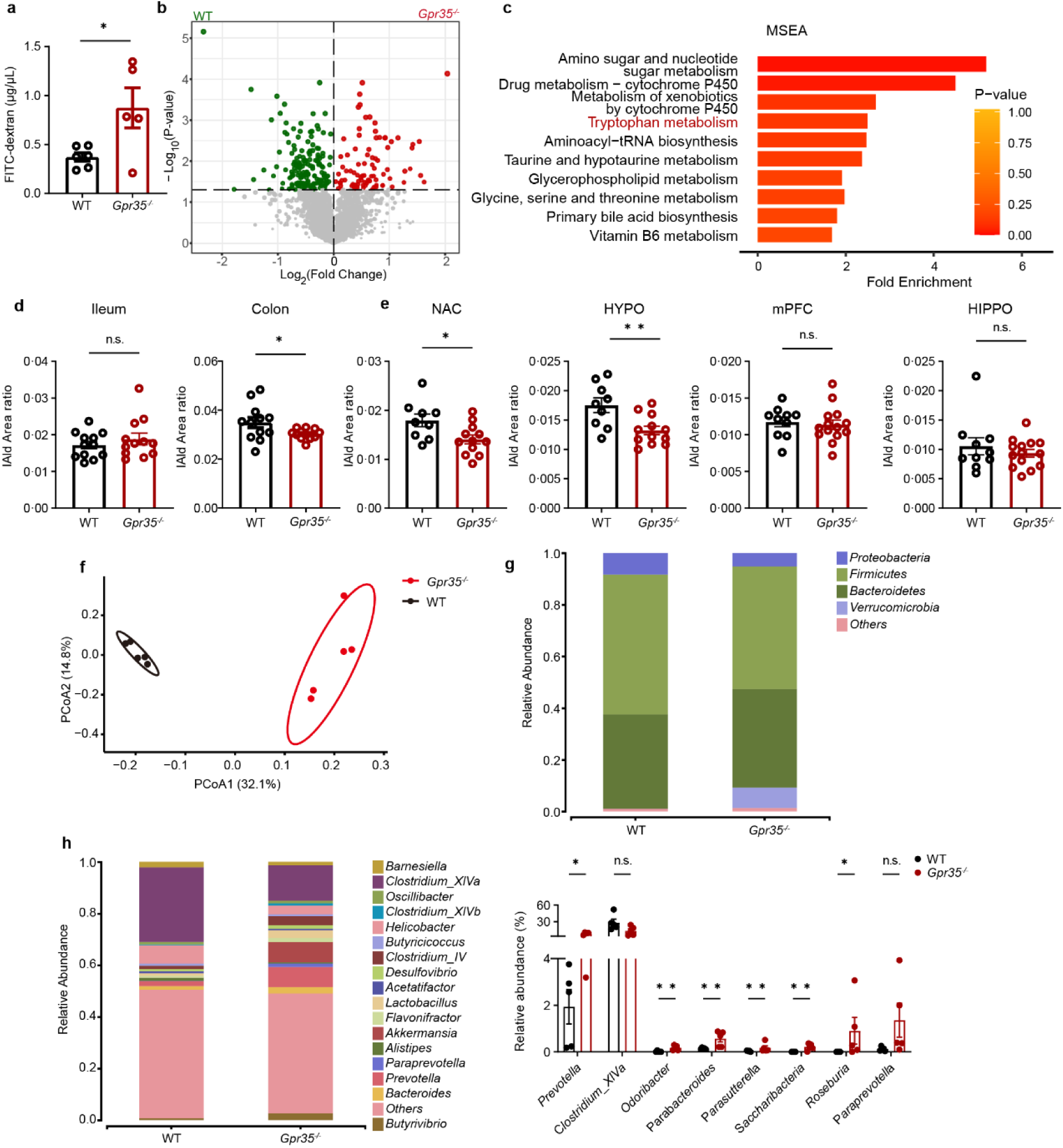
*Gpr35* ablation induces gut epithelial disturbance and microbial metabolite changes. (a) Serum concentration of FITC-dextran after oral administration to WT and *Gpr35*^-/-^ mice (n=6 and 5). (b)Volcano plotting showing differential metabolites in the serum of WT and *Gpr35*^-/-^ mice. (c)Pathway-associated metabolite set enrichment analysis (MSEA) of significantly different biological processes for *Gpr35*^-/-^ *vs* WT mice. (d, e) Relative level of IAld in the gut (ileum, colon, d) and brain (nucleus accumben, hypothalamus, hippocampus, mPFC, e) of WT and *Gpr35*^-/-^ mice (n=10-14). (f) Beta diversity of the fecal microbiome of WT and *Gpr35*^-/-^mice (n = 5) as determined by principal co-ordinates analysis (PCoA) of Bray-Curtis distances (PERMANOVA: R^2^ = 0.32, *p* = 0.008). (g,h) Averaged relative abundance of bacteria at the phylum (g) and genus (h) level. Relative abundance of differential bacteria between WT and *Gpr35*^-/-^mice at the genus level. Data represent mean ± SEM. \**p* < 0.05, \*\**p* < 0.01, n.s., no significance; two-tailed unpaired Student’s *t*-test.

Next we sought to explore gut-derived signals that may communicate with the brain to mediate the behavioral symptoms of *Gpr35*^-/-^ mice. Small-molecule metabolites from gut microorganism-host co-metabolism could signal beyond the gastrointestinal tract to modulate brain and behavior^19, 20^. Untargeted metabolomics analysis of serum enabled us to identify 263 differential metabolites, 89 of which were enriched in *Gpr35*^-/-^ mice (**Fig.2b**). Interestingly, the top four most differential pathway include tryptophan metabolism which is well known for behavioral regulation^19^ (**Fig.2c**). We confirmed a significant change in host and microbiome-derived tryptophan metabolites in the serum by targeted metabolomics (**Fig.S4a**). Following this clue, we further profiled the tryptophan metabolites in the ileum, colon tissue and several brain regions involved in behavioral regulation. We noted an extensive change of several indole metabolites, including IAld that was significantly depleted in the colon, hippocampus (HIPPO), nucleus accumbens (NAC), mPFC and hypothalamus (HYPO) (**Fig.2d,e**), as well as indole-3-acetate (IAA) that was markedly diminished in the HIPPO (**Fig.S4b**). Meanwhile, in line with the behavioral changes, we found that several neurotransmitters such as γ-aminobutyric acid (GABA) and glutamate were consistently altered in the gut and brain (**Fig.S4c**). Altogether, these findings suggest that *Gpr35* deficiency reconfigures microbial indole metabolites of tryptophan that may signal along the gut-brain axis to mediate the behavioral changes.

How did *Gpr35* deficiency impact microbial metabolism? To answer this question, we further performed *16S* ribosomal RNA (rRNA) sequencing of fecal microbiome. While we found no major changes in α-diversity (**Fig.S4d**), β-diversity analysis measured by Bray-Curtis distance revealed clustering of samples by genotype (**Fig.2f**), indicating significant changes in gut microbial composition after *Gpr35* ablation. Specifically, the relative abundance of *Lachnospiraceae* and *Clostridium_XIVa* was decreased while *Parabacteroides* and *Parasutterella* spp. were significantly expanded in *Gpr35*^-/-^ mice compared to the control littermates (**Fig.2g,h** and **Fig.S4e**). We also confirmed a similar remodeling of gut microbial structure between *Gpr35*^flox^ and *Villin-cre*; *Gpr35^flox^* mice (**Fig.S4f,g**). Together, these findings suggest a potential role of gut microbial remodeling in the depression-like behavior of *Gpr35*^-/-^ mice.

### Gut microbiome mediates the behavioral abnormality of Gpr35^-/-^ mice

Next we sought to test whether the dysbiotic microbiome elicited by *Gpr35* ablation could directly influence the brain and behavior. To this end, firstly we employed fecal microbiome transplantation (FMT) from CON or *Gpr35*^-/-^ donors into ABX-treated recipients (labeled ‘WT recipient’’ or ‘‘KO recipient’’) by oral gavage for 2 weeks (**Fig.3a**). After, bacterial *16S* rRNA gene sequencing of fecal contents from recipient mice revealed preserved alterations in the abundance of certain taxa between donor mice and their corresponding recipients (**Fig.3b** and **Fig.S5a**), including the *Clostridium_XIVa, Parabacteroides,* and *Bacteroidales* genus (**Fig.S5b**). In the series of behavioral tests, we found that the behavior of the recipient mice largely mirrored that of their respective microbiota donors especially for anxiety and social avoidance (**Fig.3c-e**), indicating a potential cause-effect relationship between the microbiota and depressive behavior.

**Fig.3.**
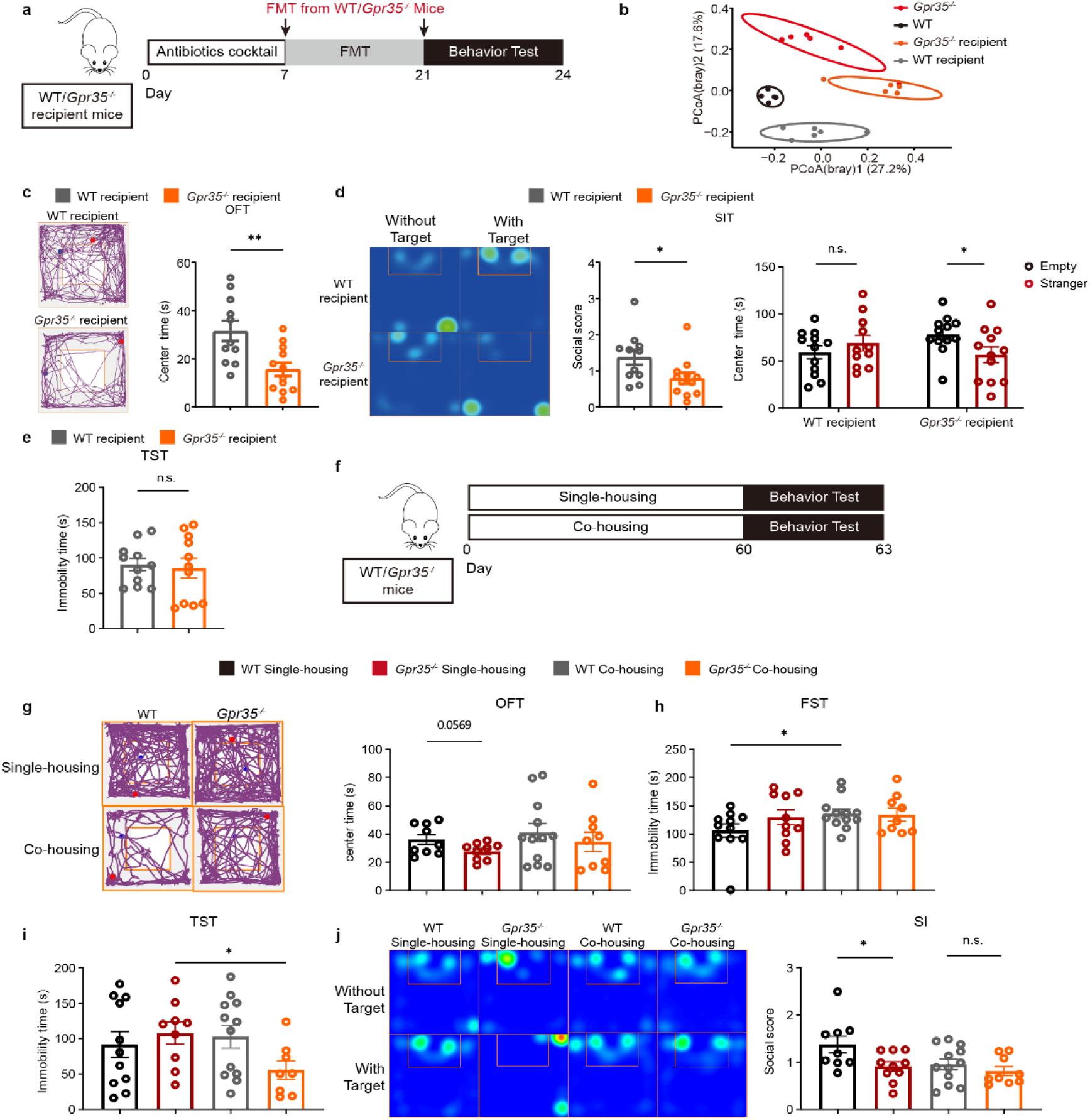
Gut dysbiosis contributes to the behavioral abnormality of *Gpr35*^-/-^ mice. (a) Experimental scheme of fecal microbiome transplantation (FMT) from WT or *Gpr35*^-/-^ donor mice. (b) PCoA plotting of fecal microbial composition of FMT donor and recipient mice, presented as averaged relative abundance at the genus level (PERMANOVA: R^2^ = 0.41, *p* = 0.00016). (c-e) Behavioral assessment of recipient mice in the open field test (c), tail suspension test (d) and social interaction test (e). (f) Experimental scheme of the co-housing experiment. (g-j) Behavioral assessment of separately and co-housed mice in the open field test (g), forced swimming test (h), tail suspension test (i) and social interaction test (j). n=8-12. Data represent mean ± SEM. \**p* < 0.05, \*\**p* < 0.01, n.s., no significance; two-tailed unpaired Student’s *t*-test.

To further ascertain the causal role, we co-housed littermate WT and *Gpr35*^-/-^ mice at the age of week 6 to induce gut microbial interchange (**Fig.3f**). After co-housing for 8 weeks, we checked the gut microbiome and behavior of WT mice, *Gpr35^-^*^/-^ mice, and co-housed counterparts, which confirmed that the gut microbial structure is transferrable under this paradigm (**Fig.S5c**). Supporting a causal role of gut microbiome, co-housed WT and *Gpr35*^-/-^ mice showed largely similar performance in most of the behavioral tests (**Fig.3g-i**). Moreover, along with the change of gut microbial configuration in separate and co-housed counterparts (**Fig.S5d,e**), we found that co-housing conferred pro-despair and anxiogenic effects on WT mice (**Fig.3h**). Taken together, FMT and cohousing experiments confirm a direct link between gut dysbiosis and depression-like behavior in *Gpr35*^-/-^ mice.

### Parabacteroides distasonis enriched in Gpr35^-/-^ mice induces depression-like behavior

To identify causal microbes from the differential microbiome between WT and *Gpr35*^-/-^ mice, we resorted to the spatiotemporal patterns of gut microbiome in the FMT and co-housing experiments. Lefse analysis revealed that *Parabacteroides, Clostridium, Phascolarctobacterium* spp. are among the several most distinct genera between the recipient mice (**Fig.S6a**). We then identified several species (e.g., *Bacteroides sartorii*, *Clotridium scindens, Parabacteroides distasonis, Parabacteroides gordonii*, *Parasutterella excrementihominis*) that exhibited consistent alterations in the donor and recipient mice (**Fig.4a**). We further looked at the microbial profile in the co-housing study, and found a notable transmission of *P.excrementihominis* and *Parabacteroides distasonis* with behavioral changes of WT counterparts (**Fig.4b**). In consideration of all these parameters, we tested 3 strains (*P. distasonis*, *P.excrementihominis, Clotridium Scindens)* for their behavioral impacts. To explore the potential pro-depressant effect, we mono-inoculated anaerobic cultures of the strains into antibiotic pre-treated WT mice, by repeated oral administration (approximately 1-3*10^8^ cfu once daily) at 2-day intervals for a total of 3 treatments (**Fig.4c**). We collected the feces of mice after the behavioral test and confirmed that the gene transcripts were significantly increased in recipient mice (**Fig.S6b**), supporting the survival of bacteria following oral gavage. Supplementation of antibiotic-treated WT mice with live *P.distasonis* or *P*. *excrementihominis* induced depression-like behaviors (**Fig.4d-h**), whereas treatment with heat-killed bacteria showed no effects on the behavioral parameters, suggesting that metabolically active bacteria are required for the behavioral changes. These behavioral changes were not accompanied by glial or neurogenesis changes in the brain (**Fig.S6c,d**). Because *P.distasonis* and *P*.*excrementihominis* are gram-negative bacteria, we further investigated the effect of selective bacteria depletion on *Gpr35* loss-associated behavior. Neomycin treatment, which depletes gram-negative bacteria, abolished the discrepancy in anxiety and depression-like behavior of *Gpr35*^-/-^ and WT mice (**Fig.S6e-g**). We also explored the potential anti-depressant effect of *Clostridium scindens* in UMS mice (**Fig.4i**). Of interest, we found that *Clotridium scindens* relieved the depression-like behavior of UMS mice (**Fig.4j-l**), suggesting that the collective changes of the three species may contribute to the behavioral changes of *Gpr35*^-/-^ mice.

**Fig.4.**
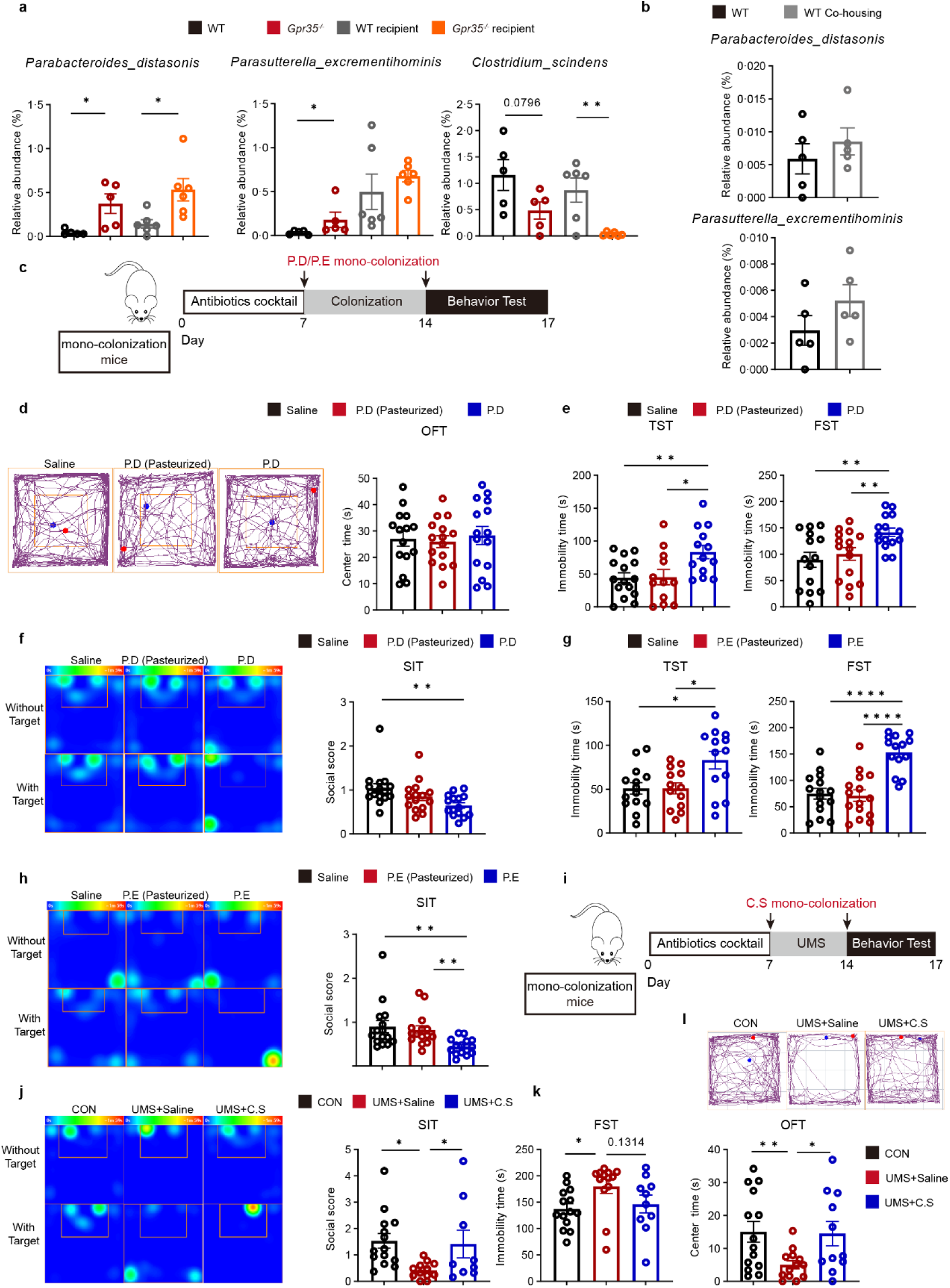
Select microbes from *Gpr35*^-/-^ mice modulate depressive behavior. (a) Relative abundance of bacterial species that are consistently changed between the WT and *Gpr35*^-/-^ donor mice and their recipient group. (b) Relative abundance of bacterial species that are altered by co-housing in WT mice. (c) Experimental scheme of *P.distasonis* (*P.D*) or *P.excrementihominis* (*P.E*) colonization to ABX-treated mice. (d-f) Behavioral assessment of *P.distasonis* colonized mice in the open field test (d), tail suspension test (e) and social interaction test (f). (g,h) Behavioral assessment of *P.excrementihominis* colonized mice in the tail suspension test (g) and social interaction test (h). (i) Experimental scheme of *Clotridium Scindens* (*C.S.*) colonization to ABX-treated mice for anti-depressant effects. (j-l) Behavioral assessment of *Clotridium Scindens* colonized mice in the social interaction test (SIT, j), forced swimming test (FST, k) and open field test (OFT, l). Data represent mean ± SEM. \**p* < 0.05, \*\**p* < 0.01, \*\*\*\**p* < 0.0001; two-tailed unpaired Student’s *t*-test.

To investigate how much the social deficits were related to metabolic changes in mice, we performed targeted metabolomic profiling of tryptophan metabolites and neurotransmitters along the gut-brain axis. Serum levels of IAld, 5-HIAA, GABA and glutamate were significantly lower in *P. distasonis* colonized mice (**Fig.S7a**). In the NAC and mPFC, we found a remarkably reduced level of IAld and glutamate, accompanied by significantly increased indole-3-lactate (ILA) (**Fig.S7b**). In line with the report of *P. distasonis* in the regulation of bile acid metabolism, and, we found that *P. distasonis* colonization affected few bile acids such as αMCA and TβMCA (**Fig.S7c**). Taken together, these results suggest a crucial role of *P. distasonis* in mediating the behavioral abnormality of *Gpr35*^-/-^ mice, possibly via rewiring gut -to-brain metabolic signals.

### Microbial IAld mediates the behavioral changes of Gpr35^-/-^ mice

Having established the role of gut dysbiosis in the behavioral change of *Gpr35*^-/-^ mice, next we sought to clarify the key microbial metabolites mediating these effects. We profiled indole metabolites in the recipient mice to seek for those featuring similar patterns of change as the donor mice. We found a remarkable decrease of IAld in the serum of mice receiving fecal microbiome from *Gpr35*^-/-^ mice (**Fig.S8a**), which was also consistent with that found in *P. distasonis* colonized mice (**Fig.S7a**). Moreover, a strong negative correlation was observed for the abundance of *P. distasonis* and serum IAld, but not IAA or ILA, in WT and *Gpr35*^-/-^ mice (**Fig.5a** and **Fig.S8b**). Moreover, we found that co-housing induced transmissible changes of IAld in the NAC and mPFC of *Gpr35*^-/-^ mice (**Fig.5b** and **Fig.S8c,d**), which indicates a strong association of this metabolite with depression-like behavior To directly determine the functional role of indole metabolites in behavioral regulation, we tested the potential protective effects of IAld by supplementing IAld to UMS-exposed WT mice (**Fig.5c**). IAld gavage brought brain IAld concentration to levels similar to non-stressed mice **(Fig.S8e**). Notably, IAld treatment effectively rescued most of the core symptoms of depression of UMS mice, as shown by the immobility time in the FST and TST and the time in the central area of OFT (**Fig.5d-f**). Considering the marked increase in ILA balance in mouse brain and intestine of *P.distasonis* colonized mice (**Fig.S7b**), we also determined the regulatory role of ILA in depressive behavior (**Fig.5g**). We found that oral gavage of ILA increased brain ILA concentration (**Fig.S8f**) and triggered a spectrum of depression-like behavior in mice, which typically include social avoidance, declined central area entrance and increased immobility (**Fig.5h-j**). Given the imbalance in IAld and ILA triggered by *P.distasonis*, these findings indicate that the diminishment of IAld in the context of ILA increase could collectively contribute to the depression-like behavior of *Gpr35*^-/-^ mice.

**Fig.5.**
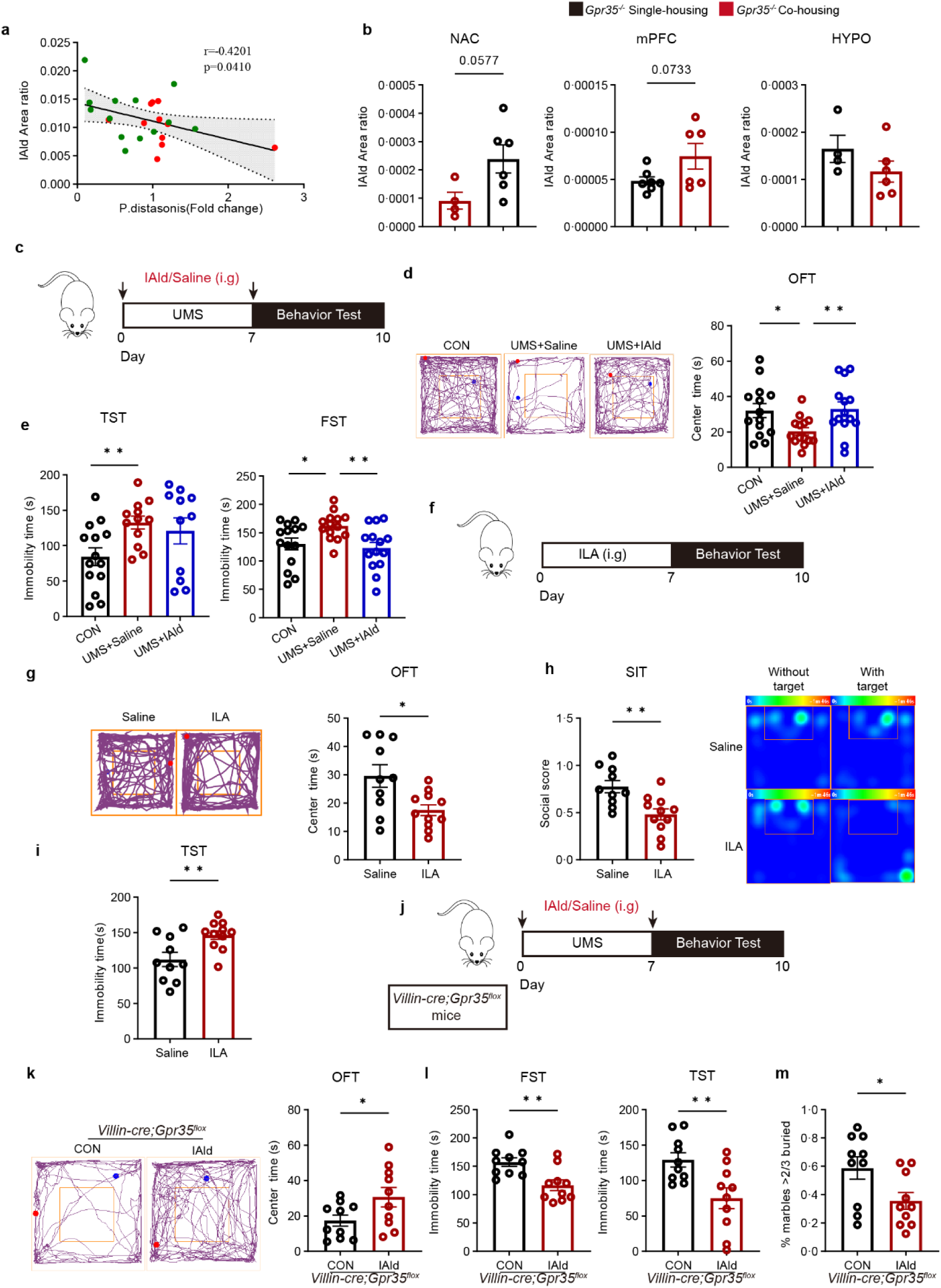
Microbial metabolites of tryptophan counteractively modulate depressive behavior. (a) Spearman correlation analysis of serum IAld and *P. distasonis* abundance in *Gpr35*^-/-^ (red) and WT (green) mice. (b) IAld in the brain regions of single and co-housed *Gpr35*^-/-^ mice. (c-e) Schematic of IAld treatment (c) and behavioral assessment of vehicle and IAld (20 mg/kg, i.g.)-treated UMS mice in the open field test (d) and tail suspension test (e). (f-i) Schematic of ILA treatment (f) and behavioral assessment of vehicle or ILA-treated WT mice in the open field test (g), tail suspension test (h) and social interaction test (i). (j) Schematic of IAld (20 mg/kg, i.g.) treatment to *Villin-cre*;*Gpr35^flox^*mice. (k-m) Behavioral assessment of vehicle and IAld-treated *Villin*-*cre*;*Gpr35^flox^* mice in the open field test (k), tail suspension test (l) and marble burying test (m). Data represent mean ± SEM. \**p* < 0.05, \*\**p* < 0.01. two-tailed unpaired Student’s *t*-test.

We also wondered whether IAld treatment would be sufficient to rescue the depression-like phenotypes in *Gpr35*^-/-^ mice. To answer this question, *Gpr35*^-/-^ mice were orally administered with IAld (20 mg kg^−1^) or vehicle every 2 days for 3 weeks and then examined for the behavioral impacts (**Fig.5k**). Notably, oral treatment with IAld significantly and robustly improved the despair and anxiety-like behavior (**Fig.5l-n**). Thus, the decrease in IAld may underlie the depressive behavior induced by Gpr35 deficiency.

### IAld and ILA counteractively regulate synaptic plasticity and depressive behavior

To investigate the molecular signatures that underlie gut microbiome-modulated susceptibility to depression, we performed RNA sequencing on representative brain regions (NAc, HIPPO, mPFC) regulated by the microbial metabolites. UMS exposure and IAld treatment each produced strong impacts on the gene expression program in the NAC and HIPPO, as indicated by differential gene expression analysis (**Fig.6a** and **Fig.S9a-d**). In particular, IAld treatment affected genes were mostly enriched to neuronal transmission and synaptic plasticity in Gene Ontology (GO) categories (**Fig.6b**). Notably, 82 gene were selectively upregulated by IAld in the NAc of UMS mice (**Fig.6c**), suggesting that IAld could promote the restoration of neuroplasticity in stressed mice brain. These findings were well aligned by RNA sequencing on representative brain regions from WT and *Gpr35*^-/-^ mice, which also indicated disrupted synaptic transmission and neurogenesis as a prominent feature (**Fig.6c,d** and **S9e,f**). We also examined the impact of ILA supplementation on gene expression in the brain. In line with a pro-depressive effect of ILA, GO enrichment analysis indicated that ILA negatively affect synaptic transmission (**Fig.6e,f** and **Fig.S9g**). To explore whether IAld could directly affect neuronal growth, we treated primary neurons with IAld. Consistent with the findings in mice, IAld treatment promoted neuron growth (**Fig.6g,h**).

**Fig.6.**
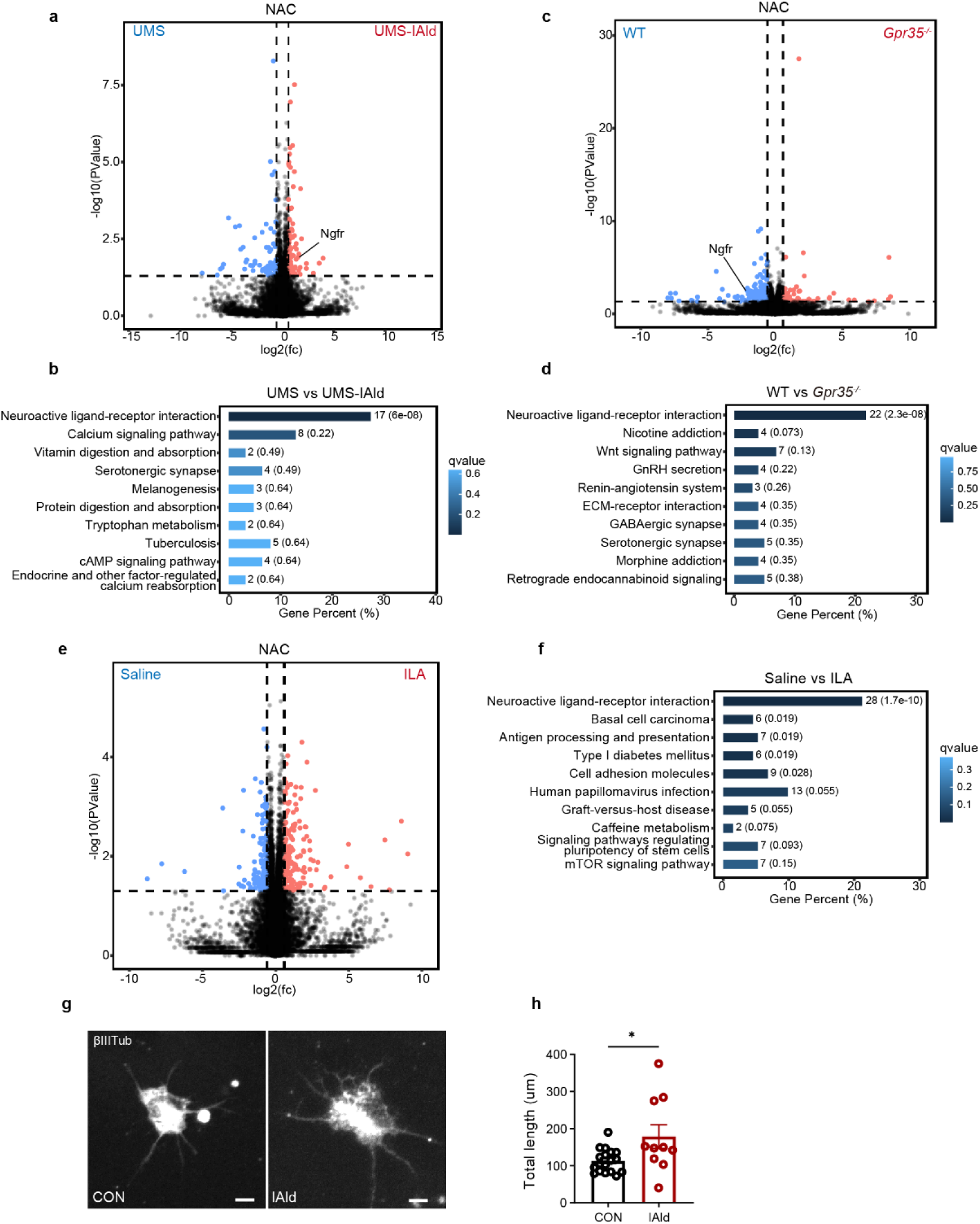
IAld potentiates neuroplasticity and rescues the behavioral abnormality of ***Gpr35*^-/-^ mice.** (a)Volcano plot displaying 158 differentially expressed genes (DEGs, log2 fold change >0.2, adjusted *p* < 0.05) in NAc from UMS *vs* UMS+IAld mice. (b)Top 10 significantly enriched KEGG pathways in NAc of UMS vs UMS+IAld group (n= 3 independent biological samples per group). (c)Volcano plot displaying 223 DEGs in NAc from WT vs *Gpr35*^-/-^ mice. (d) KEGG pathway analysis of differential biological processes in the NAc from WT vs *Gpr35*^--^ mice. n=3 mice/group. (e)Volcano plot displaying 331 DEGs in NAc from vehicle vs ILA mice. (f)Top 10 significantly enriched KEGG pathways in the NAc of vehicle vs ILA group (n= 3 mice/group). (g) Representative ßIII-Tubulin (ßIIITub) staining images of primary neurons treated with IAld (1 μM). (h) Quantification of neurite outgrowth (n=10-15 neurons per group), Scale bar: 5 µm. Data represent mean ± SEM. \**p* < 0.05. two-tailed unpaired Student’s *t*-test.

### Depressive patients feature depleted IAld and enriched P.distasonis

To gain translational insights into our findings, we performed a retrospective analysis of indole metabolites in the serum of patients with major depressive disorder (MDD, n=77) and healthy controls (HC, n=85). The serum concentration of IAld and indole-3-acetamide (IAM), a precursor of IAld, was significantly decreased in MDD patients (**Fig.7a** and **Fig.S10a**). We also examined the prevalence of *P.distasonis, P.excrementihominis* and *C. scindens* in the fecal microbiome of HC and MDD patients. The results showed that *P. distasonis* was enriched in the gut microbiome of the MDD patients (**Fig.7b** and **Fig.S10b**), although no significant change was found for *P.excrementihominis or C. scindens* (**Fig.S10c,d**). To examine whether this was true for other neuropsychiatric disorder, we also explored their differences between HC and bipolar disorder (BD) and bipolar mania (BM) patients. Of note, we observed a consistent enrichment of *P.excrementihominis* in the gut microbiome of the two patient cohorts (**Fig.7b** and **Fig.S10b**). To further explore the relationship between serum IAld and depressive behavior, correlation analyses were conducted. The result shows that the severity of depression, as assessed by the HAMD score, were negatively associated with serum IAld (**Fig.7c**), which strengthens our hypothesis that serum IAld could be functionally involved in the dysregulated gut-brain axis of depressive patients.

**Fig.7.**
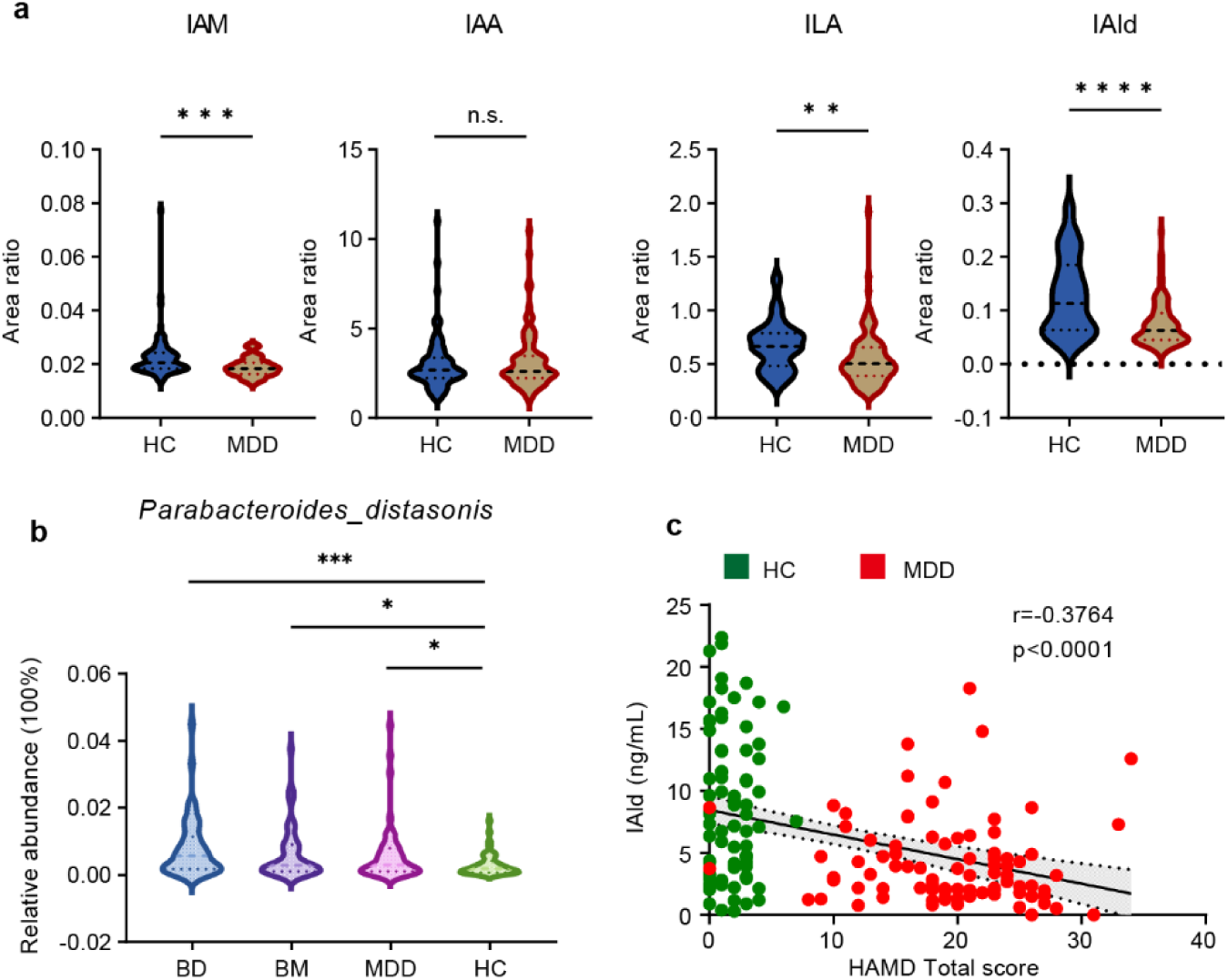
IAld is depleted in the serum of depressive patients. (a) Violin plot of the relative abundance of bacterial tryptophan metabolites in the serum of healthy control (HC, n-77) and major depressive disorder (MDD, n=85) patients. IAM, indole-3-acetamide; IAA, indole-3-acetate; ILA, indole-3-lactate; IAld, indole-3-carboxaldehyde. (b) Relative abundance of *Parabacteroides distasonis* in the gut microbiome of HC, MDD, bipolar disorder (BD) and bipolar mania (BM) patients by *16S* rRNA sequencing (n=44-51). (c) Correlation of serum IAld with the depression score of MDD patients. \**p* < 0.05, \*\**p* < 0.01, \*\*\**p* < 0.001, \*\*\*\**p*< 0.0001.

## Discussion

Despite the growing prevalence of neuropsychiatric disorder and the large efforts spent in identifying genetic basis of susceptibility, there is limited insights into the role of environment-gene interactions and mechanistic links with the behavioral symptoms. Here we show that *Gpr35*-microbe interplay dictates the susceptibility to depressive behavior. More importantly, we identify gut microbial catabolites of tryptophan as counteractive signals shaping neuronal plasticity and depressive behavior. These findings reveal mechanistic interrelationships between host genetics, the microbiota, and metabolite-mediated depressive behavior, suggesting potential diagnostic and therapeutic approaches of depressive disorders.

Mouse models of neuropsychiatric diseases are essential for mechanistic understanding and development of new therapies. Our data identify that *Gpr35* variant, which is previously established as a risk gene for inflammatory bowel disease, could induce and predispose to depressive behavior in mice. *Gpr35* knockout mice is characterized by gut epithelial, microbial disturbance and neuroplasticity impairment, without elevated neuroinflammatory manifestations associated with glial activation. Since *Gpr35* global and gut epithelia knockout mice display many core features of depressive behavior such as social avoidance, anxiety and despair, we propose that these mice could provide a new preclinical genetic model of depression. In line with our findings, a recent study also reported that intestinal activating transcription factor 4 (ATF4) deficiency induces stress-related depressive behavior in male mice^21^. These findings underscore the importance of gut-brain axis in behavioral regulation and suggest potential gut-located targets for remote behavioral regulation.

Whereas genetic variation is believed to account for 37% of MDD cases, the vast majority of the genetic risks are of unknown etiology. Here we show that the depressive phenotype of *Gpr35* knockout mice was mechanistically linked to gut microbial remodeling. Notably, we show that *P. distasonis,* which is similarly expanded in the gut microbiome of depressive patients, was able to induce depressive behavior under physiological conditions. In support of our findings, colonization of *P. distasonis* was recently reported to induce depressive behavior in a mouse model of ileitis^22^. Although *P. distasonis* was previously known for immunoregulatory effects by regulating bile acid metabolism^23, 24^, our data reveal a new link to tryptophan metabolism and susceptibility to depression. IAld and ILA are traditionally thought to be synthesized by the gram-positive bacteria such as *Bifidobacterium adolescentis*, *Lactobacillus reuteri* and *Clostridium sporogenes*. However, our results suggest a complex regulatory mode of their catabolism and other microbes deserve attention to better understand the role of microbial tryptophan metabolism in depression. Together, these findings underscore the importance of gut dysbiosis in genetic variant-associated depression susceptibility and add *P. distasonis* as a new contributor to the list of depression-inducing microbes.

Small-molecule metabolites from gut microbiome are critical in remote control of brain and behavior. Identifying neuroactive signals and pathways that mediate the behavioral effect of the gut microbiota is of critical importance for achieving effective therapeutic intervention. Our identification of IAld and ILA to counteractively modulate neuroplasticity and depressive behaviors adds to current understanding of gut microbial metabolism in behavioral control, and suggest that the composition of these analogues may underlie the interindividual differences in depression susceptibility. In further consideration of the decreased IAld in the serum of MDD patients, the IAld/ILA ratio may provide a diagnostic marker for clinical diagnosis. IAld could rescue stress-induced depression-like behaviors in mice by augmenting synaptic transmission and neuroplasticity, which provides a novel scaffold for drug discovery aiming at bolstering neurogenesis and synaptic function. At the molecular level, this was possibly attributed to the Ngfr signaling, which deserved more exploration in future studies.

It is traditionally believed that by directly impacting brain development and function, genetic variants are the primary drivers of the behavioral symptoms associated with neurological disorders^5^. Current therapeutic approaches for neuropsychiatric disorders largely aim to target the brain directly. In our study, we confirmed that gut microbial dysbiosis induced by *Gpr35* loss was sufficient to elicit most of the depressive behavior, and neomycin treatment or supplementation of microbial metabolite IAld could reverse the behavioral symptoms. Our findings, together with previous reports in autism models and a recent study on Alzheimer’s disease^8, 25, 26^, emphasize that genetics-induced behavioral abnormalities could have gut microbial origins, and modifying the gut microbiome could provide therapeutic benefits. Therefore, to effectively reduce the burden of neuropsychiatric diseases, it is imperative to consider the complex systems pathology for target discovery and make a transition from a dogmatic brain-focused approach toward a holistic conception of therapy^27^. Investigating the mechanisms through which host and microbial factors regulate complex behaviors will not only expand our understanding of neurological disorders but may also lead to new therapies.

## Acknowledgements

This work was supported by the National Natural Science Foundation of China (grant 81930109 to H.H., grants 82274009, 81973556 to X.Z.), the National Key Research and Development Program of China (grant 2021YFA1301300 to H.H.), the “Double-first” Discipline funding of China Pharmaceutical University (grant CPUQNJC22_09 to X.Z.), and CAMS Innovation Fund for Medical Sciences (grant 2021-I2M-5-011 to G.W.).

## Author Contributions

X.Z. and H. H. conceived the project and designed the study; L.S.C. and H.Q.W. carried out most of the biochemical, molecular experiments and bioinformatics analysis, and analyzed the data, with assistance from X.Y.C and Q.W.; Y.Y.Z., Z.Y., Q.Y.Y., and Y.L.H. offered help in the animal experiments and data analysis; Y.Y. supervised clinical studies and provided clinical data; G.W. was responsible for supervising the project; H. H., X.Z., and X.L.Z. contributed to discussion and data interpretation, L.S.C., H.Q.W, X.L.Z., H.P.H and X.Z. prepared the manuscript with input from all of the authors.

## Methods

### Human Participants

We recruited major depressive disorder (MDD), Bipolar disorder (BD), bipolar mania (BM) patients and healthy control (HC) subjects at the Department of Psychology and Psychiatry, Zhongda Hospital affiliated to Southeast University (Nanjing, China). The study was approved by the Ethical Committee of Zhongda Hospital (2022ZDSYLL193-P01), and informed written consent was obtained from all participants. All the patients were initially screened by the Diagnostic and Statistical Manual of Mental Disorders (Fifth Edition, DSM-5) for inclusion. Subjects were excluded with any serious somatic diseases or systemic infection, the presence of other mental disorders, history of alcohol or substance dependence, during pregnancy or lactation, or when they received any type of anti-depressants in the 2 weeks before enrollment. No participants consumed probiotics or antibiotics in the month before enrollment in the study. A cohort of 88 MDD patients and 77 HC participants were finally sampled for serum metabolite profiling.

### Animals and treatment

All animal procedures followed the ethical guidelines outlined for the Care and Use of Laboratory Animals, and all protocols were approved by the institutional animal care and use committee of China Pharmaceutical University (No. 2020-09-017). Wild-type (WT) C57BL/6J mice (SPF, male, 5-6 weeks old) were purchased from Vital River Laboratory Animal Co., Ltd (Hangzhou, China). *Gpr35*^-/-^ mice (SPF, C57BL/6J background) were purchased from BRL Medicine Inc. (Shanghai, China). *Gpr35^flox/flox^* and *Villin1-Cre* mice were purchased from Shanghai Model Organisms Center, Inc. (Shanghai, China). All these genetic model mice were maintained and crossed at the SPF facility of GemPharmatech Co., Ltd (Nanjing, China). It is noteworthy that *Gpr35^−/−^* mice were obtained from heterozygous (*Gpr35^+/−^*) breeding, as recommended for microbiome studies^28^, ruling out potential differences in maternal care and/or early life effects on the gut microbiome. Mice were housed in standardized animal rooms (12-h light: 12-h dark cycle, room temperature 22 ± 2 °C), with food and water provided *ad libitum*.

To investigate the impact of IAld on the susceptibility to depression, WT or *Gpr35*^-/-^ mice were randomly assigned into vehicle or IAA treatment groups, which received oral administration of IAld (20 mg/kg) 0.5 h before daily UMS procedure. To investigate the impact of ILA on depressive-like behavior, ILA (20 mg/kg) was applied by oral gavage for 9 consecutive days, and behavioral tests were performed on the last 3 days). To deplete gram-negative bateria, male SPF WT (n=7) and *Gpr35*^-/-^ mice (n=6) were provided with the drinking water containing freshly prepared neomycin sulfate solution (0.5 g/L) daily for 28 days. Then, the antibiotic solution was removed and replaced with drinking water for 2 days (wash-out) before the behavioral assessment.

### Unpredictable mild stress

Mice was allowed to adapt for 1 week before the initiation of stress. The UMS regimen was based on the procedures previously described with minor modifications. Briefly, the mice were subjected to various stressors in an inevitable and unpredictable way for 1 week. Stressors, as shown in **Table S1**, were applied at random times during both night and day, in order to be completely unpredictable. Behavioral assessment was performed 1 day after the end of UMS (day 8). All animals received food and water *ad libitum* in the non-stressed period. Control mice were housed under the same laboratory conditions (12-h light: 12-h dark cycle) without stress.

### Behavioral tests

All behavioral tests were performed at 6-12 weeks of age for sexually matched mice. On each experimental day, mice were brought to the testing room and allowed to habituate for at least 1 h. The apparatus was cleaned with 75% ethanol after each trial session.

#### Open-field test

Mice were gently placed in a white plastic open-field arena (40 × 40 × 40 cm) and allowed to explore freely for 8 min. The box floor was divided into central area (20 × 20 cm) and peripheral area. The position of mice was continually monitored by a video camera and the total distance and the amount of time spent in each area were analyzed using ANY-maze software (Stoelting, USA).

#### Social interaction test

The test consisted of 2 sessions per mouse. In the first session, an empty wire-mesh enclosure was placed in the social interaction area, and the test mouse was placed in the center and allowed to explore for 2.5 min. Then, the mouse was removed from the arena and returned to its home cage until session 2. Next, in the second session, a novel CD-1 mouse was placed inside the wire-mesh enclosure, and the same test mouse was placed in the interaction area and allowed to explore for 2.5 min. The amount of time spent in the interaction zone during the absence and presence of the CD-1 mice were analyzed and social score was calculated by t2(time in Interaction area in stage2)/t1(time in Interaction area in stage1).

#### Tail suspension test

The mice were suspended by a tape attached to their tail to a horizontal hook 50 cm above the ground surface. Mouse behaviour was recorded for 6 min, and analysis of immobility time was assessed during the last 4 min. Absence of escape-oriented behavior (i.e., hanging passively, motionless) was considered immobility.

#### Forced swimming test

A water beaker was filled with 15 cm of warm water (24 ± 1◦C) arranged fresh immediately before each experiment. The mouse was placed carefully into the water and recorded for 6 min. Immobility time was assessed during the last 4 min. Animals were considered immobile when displaying no signs of escape-oriented swimming. If the animal showed any signs of distress, the test was immediately interrupted. After the test, the mice were dried with a towel and placed in a warm, new home cage.

#### Marble burying test

A standard mouse rearing cage was filled with approximately 5 cm deep clean corncob bedding material that was evenly distributed into a flat surface across the whole cage. Sixteen glass marbles (1.6 cm in diameter, plain dark glass) were then spaced evenly in a 4 × 4 grid on the surface of the bedding. During the testing phase, each mouse was placed in the cage and allowed to explore it for 30 min. At the end of the test, mice were removed from the cage and the number of marbles buried with bedding up to 2/3 of their depth was counted. Ensure consistent light source intensity and quiet test space during testing. All analyses were performed blind.

### Bio-sample collection

Following the behavioral tests, mice were sacrificed by cervical dislocation under isoflurane anesthesia. The peripheral blood was collected and the ileum and distal colon were carefully dissected and immersed in cold PBS to wash and remove fecal contents, then quickly frozen in dry ice. The brain was isolated and the hippocampus, nucleus accumbens, hypothalamus and medial prefrontal cortex were microdissected out on a cold plate and immediately frozen on dry ice. All samples were then stored at −80°C until further processing. For gut microbiome profiling, fresh fecal pellets were collected and snap frozen before analysis.

### Quantitative real-time PCR

RNA extraction was performed using RNA-easy Isolation Reagent (Vazyme) according to the manufacturer instructions. cDNA was generated using 5× HiScript III qRT SuperMix (Vazyme) according to manufacturer’s instructions. qPCR was performed using the SYBR Green Supermix Kit (Vazyme) on the CFX96 Real-Time PCR System (Bio-Rad, USA). The expression of target genes was normalized to expression of *Gapdh*, and shown as fold change relative to the control group based on the 2^-△△Ct^ method. The primer sequences are shown in **Table S2** and **S3**.

### 16S rRNA sequencing and data analysis

Library preparation, sequencing, and operational taxonomic unit (OTU) table generation were performed by BGI Genomics at Wuhan, China. Briefly, bacterial DNA was extracted using magnetic-bead based protocol. DNA quality was determined by agarose gel electrophoresis. The *16S* rDNA V3-V4 region was amplified by PCR and sequenced in the HiSeq platform (Illumina). The raw reads were processed and clustered into Operational Taxonomic Units (OTUs) using USEARCH v7.0.1090. *16S* rRNA gene sequences were clustered into OTUs at a similarity cutoff value of 97% using the UPARSE algorithm.

Principal co-ordinates analysis (PCoA) by Bray-Curtis distance was performed to visually evaluate the gut microbial difference. LDA effect size (LEfSe) was used for the identification of differentially expressed bacterial taxa between groups.

### Fecal Microbiome Transplantation

Six-week-old SPF C57BL/6J mice were administered with an antibiotic cocktail in the drinking water for 10 days. The antibiotic combination consisted of vancomycin (250 mg/L, Aladdin), metronidazole (500 mg/L, Aladdin), neomycin sulfate (500 mg/L, Aladdin) and ampicillin (500 mg/L, Aladdin). Antibiotic treated mice were maintained in sterile cages and provided sterile food and water during the course of the experiments.

On the day of transplantation, fresh fecal pellets were collected from littermate male WT and *Gpr35^-/-^* donor mice and quickly homogenized in sterile cold PBS in an anerobic chamber. After shaking for 3 min, the resulting slurry was settled for 5min and then the supernatant was isolated for FMT. Recipient mice pretreated with antibiotics as described above were then immediately colonized by a single gavage with 0.2 mL suspension every second day. Fecal samples were collected from the recipient mice 8 days later to verify colonization. Fecal samples were stored at −80 °C until further sequencing. Behavioral experiments were initiated at 6 days post-transplantation.

### Culture and colonization of bacterial strains

*Parabacteroides distasonis* (*P.distasonis,* ATCC 8503) was cultured anaerobically in ATCC Medium 1490 in an 85% N2, 5% CO2, 10% H2. To prepare control bacteria, *P.distasonis* was heat-killed by keeping the bacteria at 121°C for 120 min. Before colonization, cultures were collected and resuspended in pre-reduced sterile PBS solution. PBS, live or heat-killed *P.distasonis* were then gavaged to mice every second day for 10 days. Behavioral assays were assessed 1 day after the last time of *P.distasonis* gavage. *P. excrementihominis* (DSM 21040) were cultured at 37°C on DSMZ Medium 78, supplemented with Vitamin K1 Solution and Hematin chloride solution. *Clostridium scindens* (ATCC 35704) were cultured at 37℃ on ATCC Medium 2107. Before colonization, cultures were collected and resuspended in pre-reduced sterile PBS solution. The colonization formula was the same as above. Bacterial colonization was confirmed by qPCR amplification with primer sequences listed in **Table S4**.

### Immunofluorescence staining

Brains samples were fixed in 4% PFA at 4 °C overnight, then dehydrated in a gradient of 10%, 20% and 30% sucrose in PBS. Each dehydration was done strictly until the tissue sank to the bottom of the container. Coronal slices (12 μm) thick were obtained from frozen tissue using a sliding blade microtome then transferred to ice cold PBS. Slices were washed three times with PBS. Slices were blocked with 10% normal donkey serum, 0.3% Triton X-100 and 90%PBS (Blocking Buffer) for 1 hr at room temperature and then incubated in primary antibodies (rabbit anti-Iba1, Abcam, ab178847, 1:500; rabbit anti-GFAP, Abcam, ab7260, 1:500; rabbit anti-DCX, Proteintech, 13925-1-AP, 1:200) diluted in blocking buffer at 4°C for 24 h. For neuron dendrite staining in primary neurons, a mouse anti-beta 3 tubulin antibody (Invitrogen, MA1-118,1:200) was used.

Slices were then washed five times with 0.1% Tween-20 in PBS. Primary antibodies were visualized using secondary donkey anti-rabbit IgG H&L antibody (Alexa Fluor® 647, ab150075, 1:500) diluted in Blocking Buffer. Slices were incubated in secondary antibodies in the dark for 1h at RT. Slices were then washed five times with 0.1% Tween-20 in PBS, Nuclei were visualized with DAPI (Abcam, ab104139). Fluorescent imaging and data acquisition was performed on FV3000 (Olympus, Japan). Images were captured using Olympus SW software (Olympus, Japan). Image processing was applied uniformly across all images within a given dataset. Hippocampal DCX-expressing neuronal population and GFAP^+^ or Iba1^+^ cell number was assessed in ImageJ by an experimenter blind to group allocation.

### Histopathological analysis of intestine and colon

For hematoxylin & eosin (H&E) staining and Alcian blue (AB) staining, the distal ileum tissue was dissected and fixed in 10% formalin overnight. For H&E staining, the tissue was then embedded in paraffin and sliced into 5 mm sections, which were sequentially performed following standard procedures. For AB staining, 3% acetic acid was applied to the slides for 3 mins. AB solution (pH 2.5) was added to slides for 10 mins and then 3% acetic acid was applied to the slides for approximately 30 s to remove excess AB solution. Slides were rinsed in running tap water and then dehydrated with an ascending ethanol gradient (70%, 95%, and 100%). Slides were mounted and images were acquired under bright field in a Leica DMI 3000B light microscope (Leica, Germany) in a blinded manner.

### Intestinal permeability analysis

Briefly, mice were fasted for 12 h and then FITC-dextran 4000 (400 mg/kg body weight, 200 µL) was administered by gavage. Four hours after gavage, whole blood was collected, and serum was isolated and diluted in five-fold volume of PBS. Standard curves for calculating the FITC-dextran concentration were obtained by diluting FITC-dextran in PBS. An amount of 100 µL of both diluted animal samples and standards, as well as blank controls (PBS and diluted plasma from untreated animals) were transferred to black 96-well microplates. Analysis for the FITC-D4000 concentration was carried out with Synergy H1 reader (Bio-Tek, USA) at an excitation wavelength of 485 nm and an emission wavelength of 535 nm.

### Non-targeted metabolomics assay

The serum sample stored at −80 °C refrigerator was thawed on ice and vortexed for 10 s. 50 μL of sample and 300 μL of extraction solution (ACN: Methanol = 1:4, V/V) containing internal standards were added into a 2mL microcentrifuge tube. The sample was vortexed for 3 min and then centrifuged at 12,000 rpm for 10 min (4°C). 200 μL of the supernatant was collected and then centrifuged at 12,000 rpm for 3 min (4 °C). A 180 μL aliquots of supernatant were transferred for LC-MS analysis.

The quantitation method was developed on a Shimadzu LC20 HPLC coupled with an AB SCIEX Triple TOF-6600 mass spectrometer. Mobile phase A was 0.1% formic acid in purified water and mobile phase B was 0.1% formic acid in acetonitrile. The analysis was performed on a Waters Acquity UPLC HSS T_3_ Column(2.1×100 mm, 1.8 μm). The column temperature was 40 °C, the flow rate was 0.4 mL/min, and the injection volume was 2 μL. The flow rate was set at 0.4 mL/min with an initial mobile-phase composition of 95:5 (A:B). The mobile phase composition was raised to 10:90 (A:B) in 11 min to elute the analytes and was kept for 1 min. Finally, the composition was decreased to 95:5 (A:B) in 12.1 min and the column wash was equilibrated for 2 min. The mass spectrometer was set to positive electrospray ionization (ESI) mode or negative ESI mode, respectively. The original data file acquired by LC-MS was converted into mzML format by ProteoWizard software. Peak extraction, peak alignment and retention time correction were performed by XCMS program. The “SVR” method was used to correct the peak area. The peaks with detection rate lower than 50 % in each group of samples were discarded. After that, metabolic identification information was obtained by searching in-house standard database, integrated public database, AI database and metDNA.

### Targeted metabolomics

#### Tryptophan and neurotransmitters

Serum, brain and intestinal samples were prepared by protein precipitation, and 1-methyl-D-tryptophan(1-MT) were used as internal standard. The method was developed on a Shimadzu HPLC coupled with an AB SCIEX 5500+ mass spectrometer. Mobile phase A was 0.1% formic acid in purified water and mobile phase B was acetonitrile. The separation was performed on a Waters Atlantis T_3_ Column (2.1×100 mm, 3 μm) with the column temperature at 40 °C and the flow rate at 0.3 mL/min. The mass spectrometer was set to positive electrospray ionization (ESI) mode with selective reaction monitoring (SRM). Spray voltage, curtain gas, collision gas, ion source gas 1, ion source gas 2 and temperature were set at 5500 V, 20 psi, 10 Pa, 55 psi, 50 psi and 550 °C, respectively. Specific mass transition detection ion information was listed in **Table S5**.

#### Bile acids

Serum, brain and intestinal samples were prepared by protein precipitation. Dehydrocholic acid (dhCA) was used as internal standard. The method was developed on a Shimadzu HPLC coupled with an AB SCIEX 5500+ mass spectrometer. Mobile phase A was 0.01% formic acid in purified water and mobile phase B was 95%acetonitrile, 5% methanol and 0.01% formic acid. The analysis was performed on a Waters BEH C_18_ Column (2.1×150 mm, 1.7 μm) with column temperature at 40 °C and flow rate at 0.2 mL/min. The mass spectrometer was set to negative electrospray ionization (ESI) mode with multiple reaction monitoring (MRM). Spray voltage, curtain gas, collision gas, ion source gas 1, ion source gas 2 and temperature were set at 5500 V, 30 psi, 10 Pa, 50 psi, 50 psi and 500 °C, respectively. Specific mass transition detection ion information was listed in **Table S6**.

### Statistical analyses

Data are presented as mean ± standard error of the mean (SEM) unless otherwise described. Statistical analyses were performed with GraphPad Prism (version 9.0). Image J (version 2.3.0) was used for quantitative comparison of positive signals in immunofluorescent staining images. ImageGP, an online data visualization web server (https://doi.org/10.1002/imt2.5.), was used for plotting of heatmaps and PCoA images. Student’s *t*-test (normal distribution) or Mann-Whitney U-tests (non-normal distribution) was applied for analysis between two groups. For multiple comparisons, one-way ANOVA was used followed by Tukey’s post hoc test, as specifically described in the figure legends. No methods were used to predetermine sample size. Mice were assigned to groups randomly. Analyses were not blinded, and no datapoints were removed from the analyses. P < 0.05 was considered statistically significant.

## Data availability

The *16S* rRNA sequencing, metabolomics and transcriptomics data were deposited to China National Microbiology Data Center (NMDC), under accession numbers NMDC40038271 and NMDC40038272, respectively).

