## Supplementary Figures and Tables for "A *Gpr35* tuned gut-brain metabolic axis regulates depressive-like behavior"

Lingsha Cheng *et al.*,

**Supplementary Figs 1-10**

**Supplementary Tables 1-6**

### Supplementary Figures

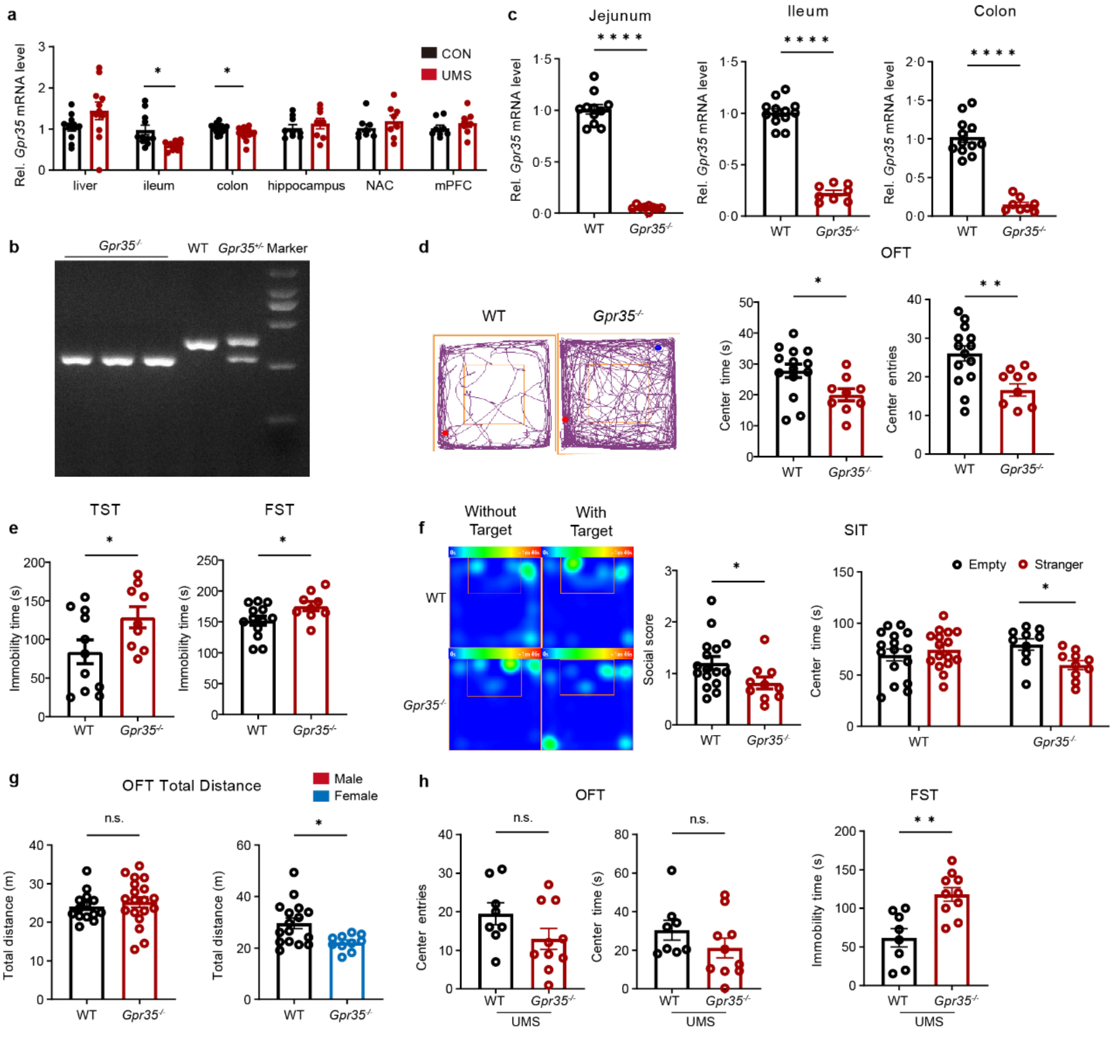


**Fig.S1** ***Gpr35*** **deficient mice exhibit depressive behavior.**

(a)*Gpr35* gene transcription levels in the liver, ileum, colon, hippocampus, nucleus accumbens (NAC) and medial prefrontal cortex (mPFC) of WT and UMS mice (n=10). (b)Representative nucleic acid electrophoresis images for the genetic identification of *Gpr35*$^{-/-}$ mice. WT: 393bp, *Gpr35*$^{-/-}$:289bp. (c)*Gpr35* gene transcription levels in the jejunum, ileum and colon of WT and *Gpr35*$^{-/-}$ mice. (d) Anxiety-like behavior of female WT and *Gpr35*$^{-/-}$ littermates as assessed in the open-field test (OFT). Time spent in the center and number of entries in central area were compared for n=16 and 10 mice. (e) Despair behavior of female WT and *Gpr35*$^{-/-}$ littermates as assessed by immobility time in the tail suspension test (TST) and forced swimming test (FST) (n = 16 and 10). (f) Social interaction score of female WT and *Gpr35*$^{-/-}$ mice as assessed in the social interaction test (SIT) (n = 16 and 10). (g) Locomotive activity of male/female WT and *Gpr35*$^{-/-}$ littermates as assessed by total distance travelled during the open-field test (OFT). (h) Anxiety and despair-like behavior of WT and *Gpr35*$^{-/-}$ littermates in the OFT and FST after exposure to UMS. Data represent mean ± SEM. *$p$ < 0.05, **$p$ < 0.01, ****$p$ < 0.0001, n.s., no significance; two-tailed unpaired Student's *t*-test.

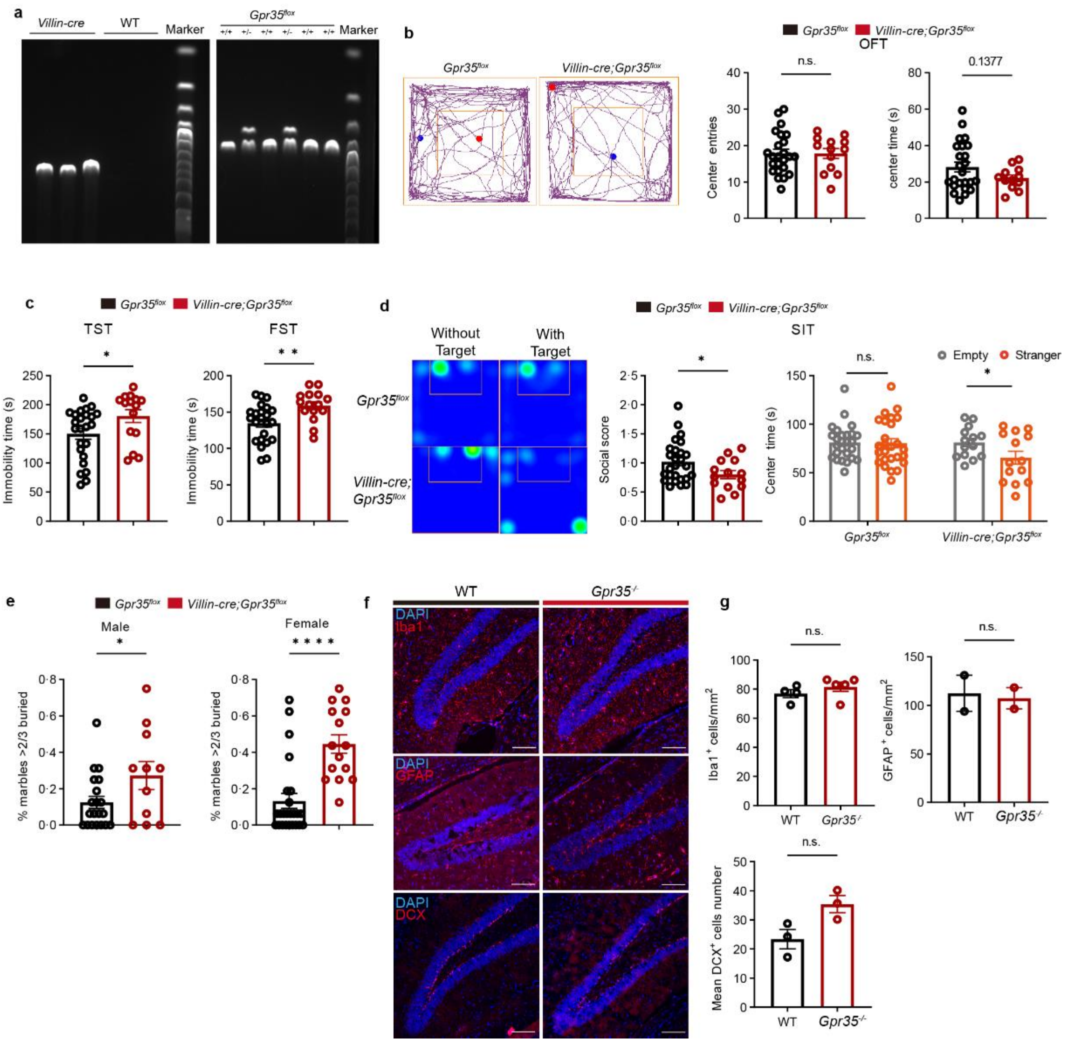


**Fig.S2 Gut epithelial specific ablation of *Gpr35* induces depressive behavior.**

(a) Representative nucleic acid electrophoresis images for the verification of conditional *Gpr35* knockout. (b-d) Depression-like behavior of female *Gpr35*$^{\text{flox}}$ and *Villin-cre*;*Gpr35*$^{flox}$ littermates as assessed in the OFT (b), TST and FST (c) and SIT (d). n = 26 and 15. (e) Anxiety-like behavior of male/female *Gpr35*$^{\text{flox}}$ and *Villin-cre*;*Gpr35*$^{flox}$ littermates as assessed in the marble-burying test. (f,g) Representative immunofluorescence images of Iba-1, GFAP and DCX from the hippocampal dentate gyrus sections from WT and *Gpr35*$^{-/-}$ mice co-labelled with DAPI (blue). Scale bar: 100 μm. Positive cells per mm$^2$ were compared for 3-5 mice. Data represent mean ± SEM. *$p < 0.05$, **$p < 0.01$, ****$p < 0.0001$, n.s., no significance; two-tailed unpaired Student's *t*-test.

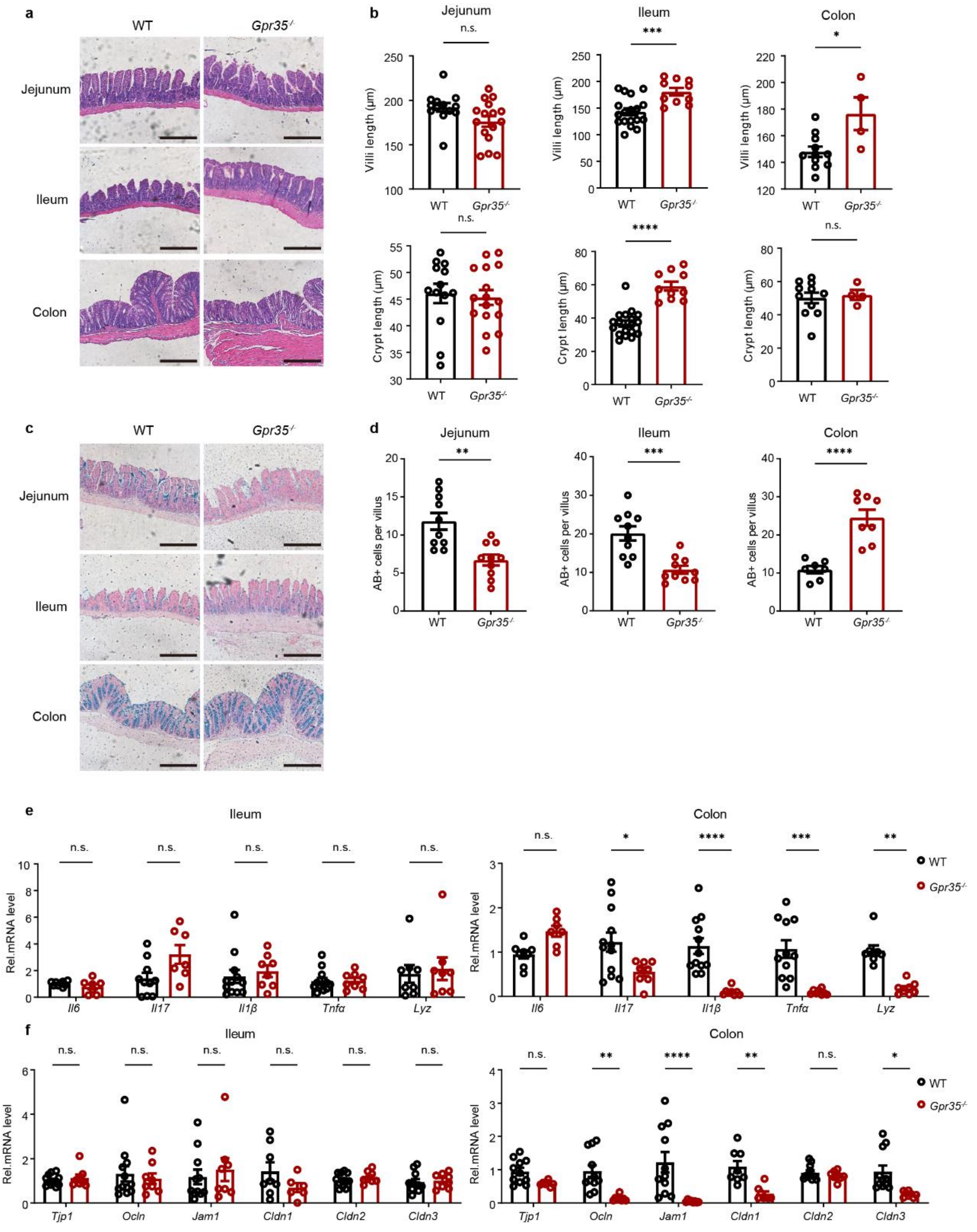


**Fig.S3 *Gpr35*$^{-}$ deficiency induces gut epithelial changes.**

(a,b) Representative H&E staining images of jejunum, ileum and colon. Scale bar: 100 μm. (c,d) Representative Alcian blue (AB) staining images of goblet cells in the jejunum, ileum and colon. Scale bar: 100 μm. (e) Relative mRNA expression of cytokines detected in the ileum (left) and colon (right) of WT and *Gpr35*$^{-/}$ mice. n=10-12. (f) Relative mRNA expressions of tight junction molecules detected in the ileum (left) and colon(right) of WT and *Gpr35*$^{-/-}$ mice. n=10-12. Data represent mean ± SEM. *$p < 0.05$, **$p < 0.01$, ****$p <$

0.0001, n.s., no significance; two-tailed unpaired Student's *t*-test. Data represent mean ± SEM. $*p < 0.05$, $**p < 0.01$, $***p < 0.001$, $****p < 0.0001$, n.s., no significance; two-tailed unpaired Student's *t*-test.

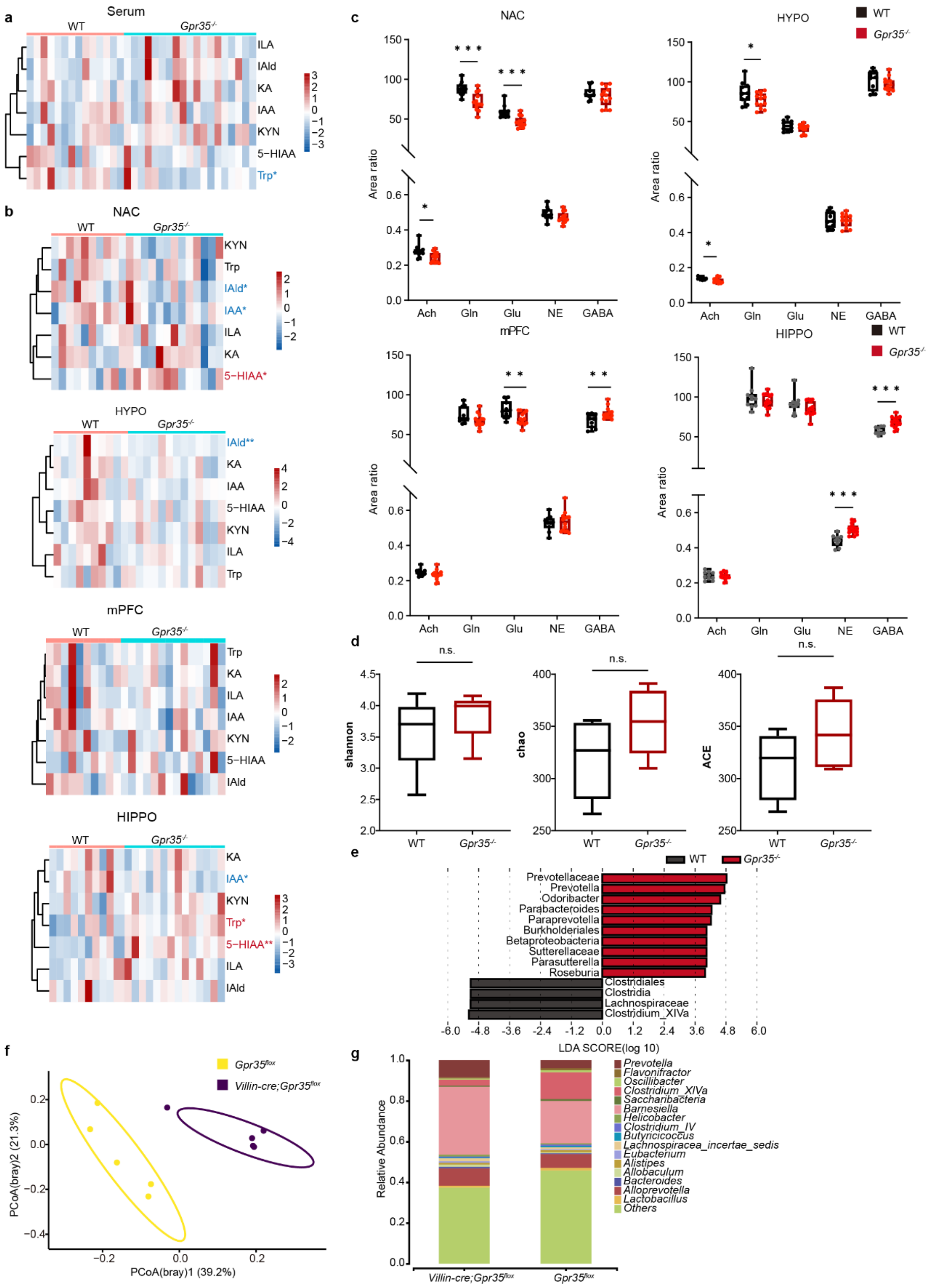


**Fig.S4 Alteration of microbial metabolite signals by *Gpr35*$^{-}$deficiency.**

(a,b) Heatmap showing tryptophan pathway metabolites in the serum (a, n=14 and 19) and nucleus accumbent (NAC), hypothalamus (HYPO), hippocampus (HIPPO) and medial prefrontal cortex (mPFC, b, n=10 and 13) of WT and *Gpr35*$^{-/-}$ mice. (c) Relative level of neurotransmitters in the NAC, HYPO, HIPPO and mPFC of WT and *Gpr35*$^{-/-}$ mice. Ach, acetylcholine; Gln, glutamine; Glu, glutamate; NE, norepinephrine; GABA, γ-aminobutyric acid. (d) Box-plots showing the α-diversity of fecal microbiota from male WT and *Gpr35*$^{-/-}$ mice, which were measured by shannon index, Chao1 and abundance-based coverage estimator (ACE). (e) Linear discriminative analysis (LDA) score of differentially expressed bacteria obtained from LEfSe analysis of the genera between the WT and *Gpr35*$^{-/-}$ mice. (f) Beta diversity of the fecal microbiome of *Gpr35*$^{flox}$ and *Villin-cre*;*Gpr35*$^{flox}$ mice (n=5) as determined by principal co-ordinates analysis (PCoA) of Bray-Curtis distances (PERMANOVA: $R^2$ = 0.34, $p$ = 0.0096). (g) Averaged relative abundance of bacteria at the genus level of the fecal microbiome of *Gpr35*$^{flox}$ and *Villin-cre*;*Gpr35*$^{flox}$ $^{-}$mice. Data represent mean ± SEM. *$p$ < 0.05, **$p$ < 0.01, ***$p$ < 0.001, n.s., no significance; two-tailed unpaired Student's *t*-test.

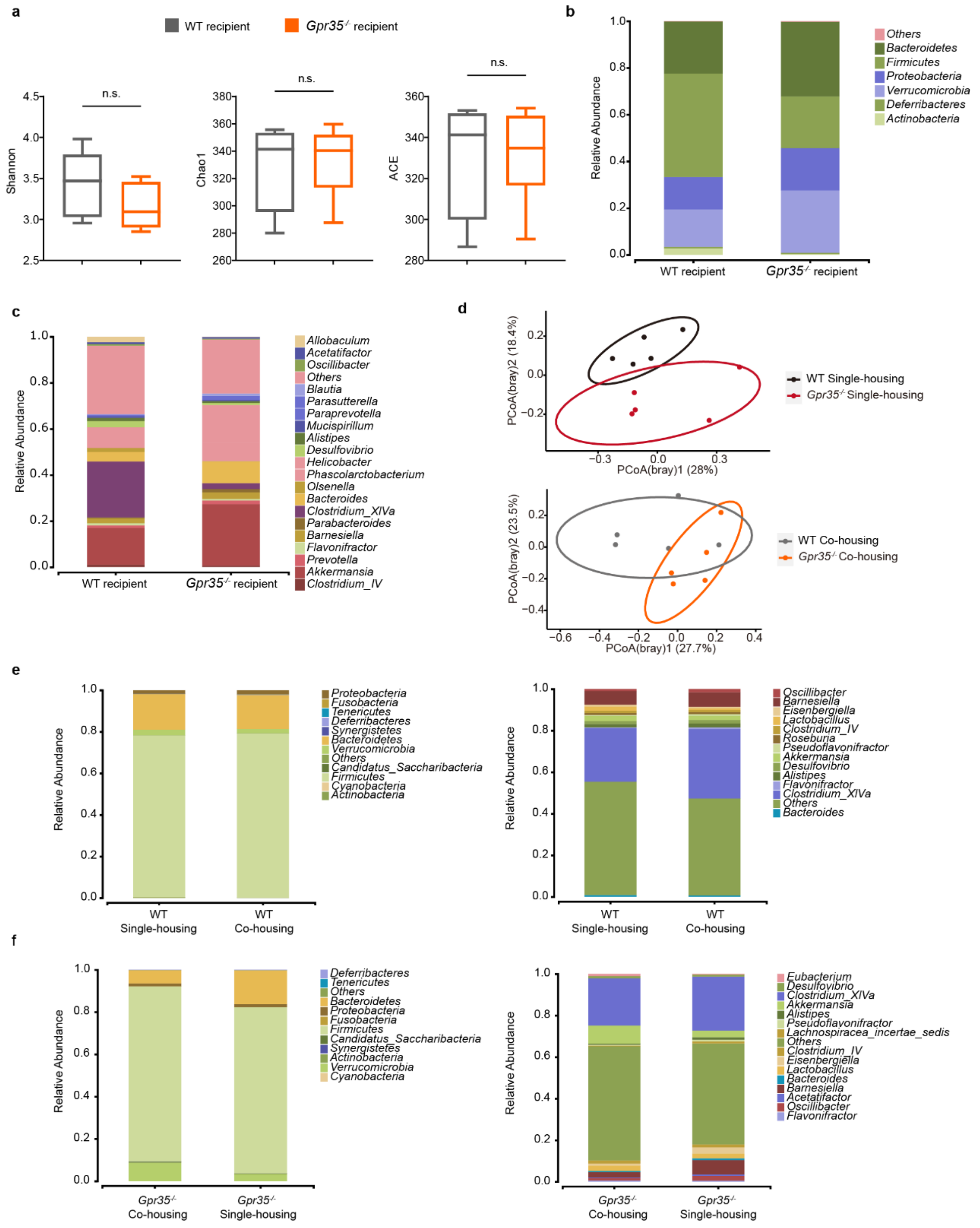

**Fig.S5. Gut microbiome transmits the behavioral phenotype of *Gpr35*$^{-/-}$ mice.**

(a) Box-plots showing the α-diversity of fecal microbiome from the recipient mice, which were measured by shannon index, Chao1 and abundance-based coverage estimator (ACE). (b, c) Averaged relative abundance of bacteria at the phylum (b) and genus (c) level in the recipient mice. (d) PCoA plotting of fecal microbial composition of separately (left; PERMANOVA, $R^2$ = 0.18, $p$ = 0.028) or co-housed (right, $R^2$ = 0.17, $p$ = 0.085) WT and *Gpr35*$^{-/-}$mice (n=5). (e,f) Average relative abundance of bacteria in WT (e) or *Gpr35*$^{-/-}$(f) mice under separate or co-housing conditions. n.s., no significance.

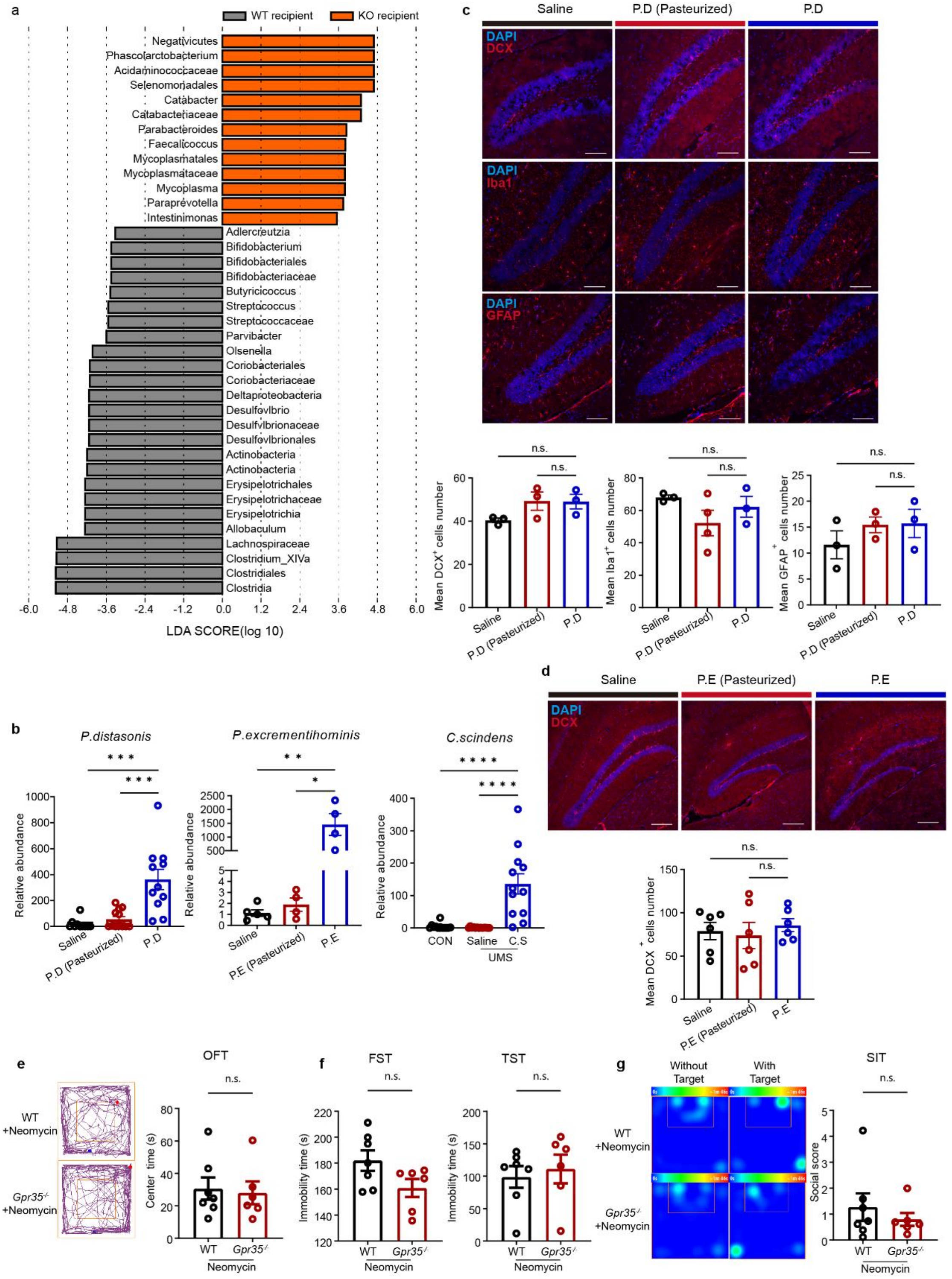


**Fig.S6 *P.distasonis* colonization induces behavioral abnormality reminiscent of *Gpr35*$^{-/-}$ mice.**

(a) LEfSe analysis of the differentially enriched genera between the recipient mice of WT and *Gpr35*$^{-/-}$ donors. (b) Verification of *P.distasonis*, *P.excrementihominis, C.Scindens* colonization by PCR. (c) Representative immunofluorescence images of Iba-1, GFAP and DCX from the hippocampal dentate gyrus sections of *P.distasonis* colonized mice. Scale bar:100μm. (d) Representative immunofluorescence images of DCX from the hippocampal dentate gyrus sections of *P.excrementihominis* colonized mice. Scale bar: 100 μm. (e-g) Behavioral assessment of neomycin-treated WT and *Gpr35*$^{-/-}$ mice in the open field test (e), tail suspension test (f) and social interaction test (g). Data represent mean ± SEM. **p* < 0.05, ***p* < 0.01, ****p* < 0.001, *****p* < 0.0001; n.s., no significance. two-tailed unpaired Student's *t*-test.

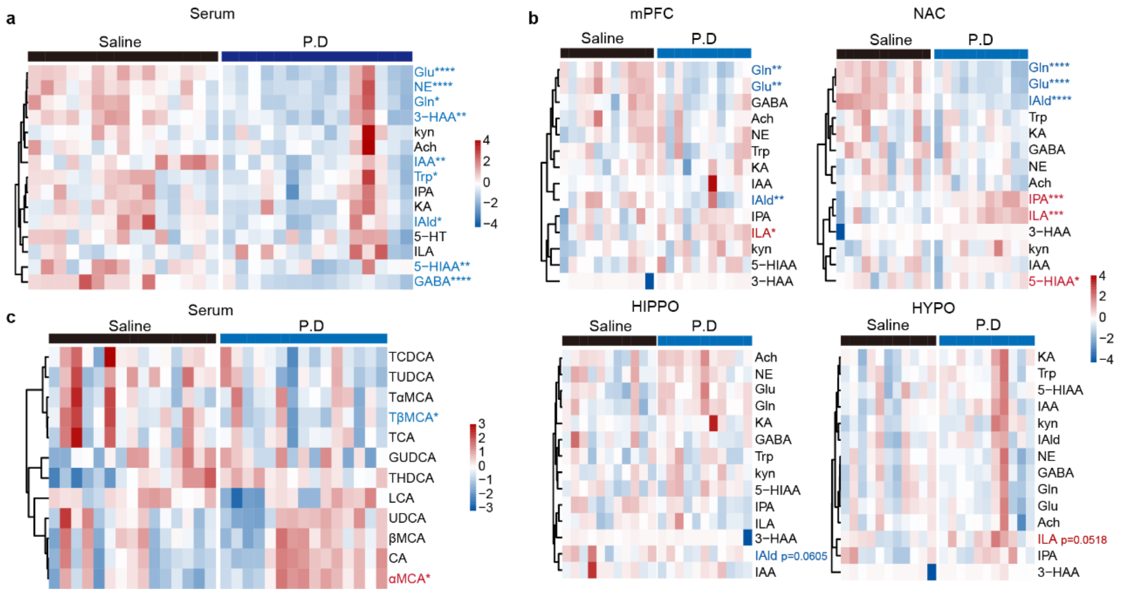


**Fig.S7 *P.distasonis* colonization induces metabolic changes in the serum and brain.** (a) Tryptophan pathway metabolites in the serum of *P.distasonis* colonized mice. (b) Tryptophan pathway metabolites in the brain of *P.distasonis* colonized mice. (c) Bile acid metabolites in the serum in *P. distasonis* colonized mice. *$p < 0.05$, **$p < 0.01$, ***$p < 0.001$, ****$p < 0.0001$; two-tailed unpaired Student's *t*-test.

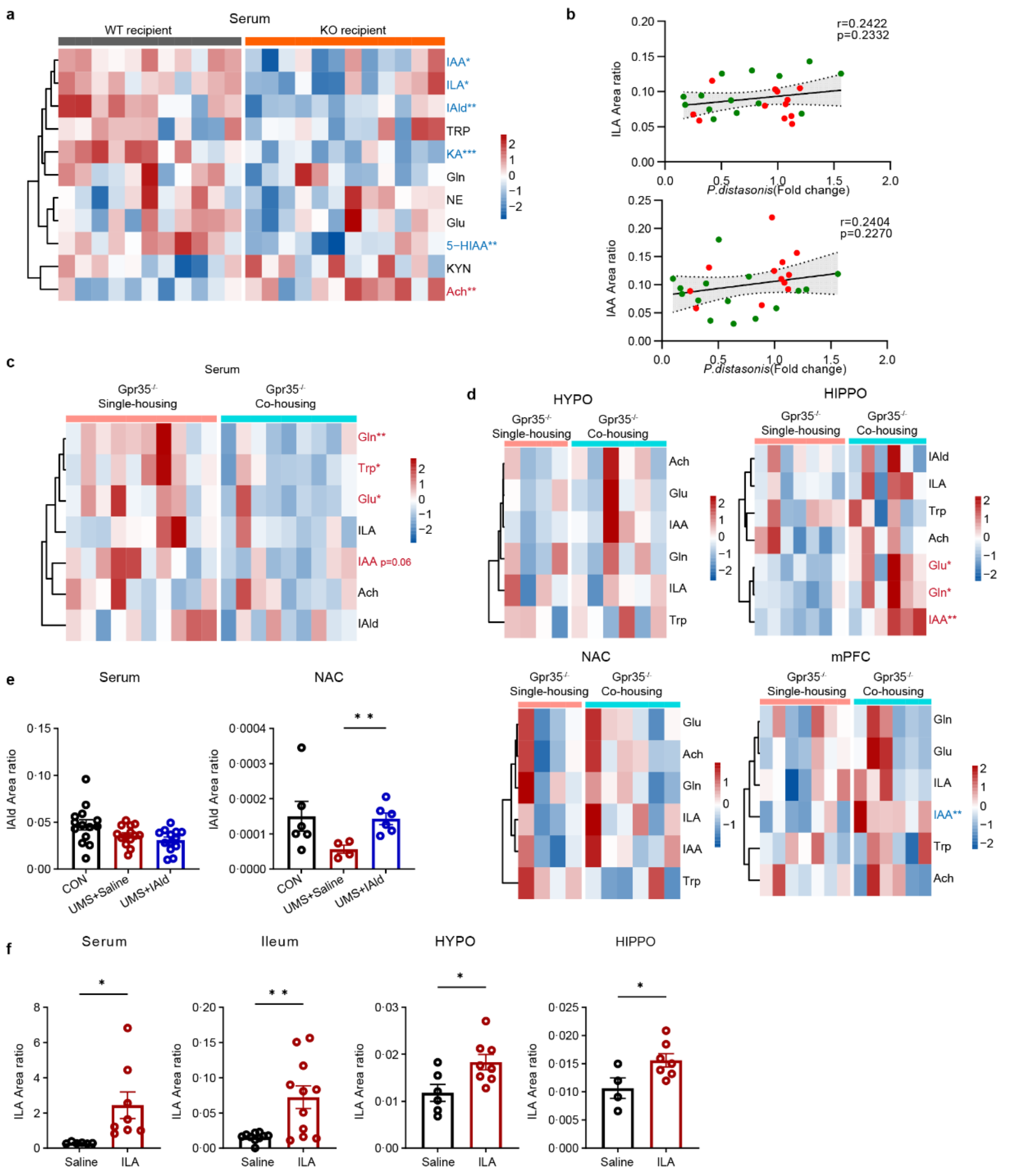


**Fig.S8 Association between IAld and ILA with depressive behavior.**

(a)Tryptophan pathway metabolites in the serum of recipient mice (n=11-12). (b) Spearman correlation analysis of serum ILA and IAA with *P. distasonis* abundance in *Gpr35*$^{-/-}$ (red) and WT (green) mice. (c,d)Tryptophan pathway metabolites in the serum (c, n=9-10) and brain regions (d, n=4-7) of single or co-housed *Gpr35*$^{-/-}$mice. (e) Serum and brain level of IAld after its oral supplementation to UMS mice (n=13-14 and 4-6). (f) Peripheral and brain level of ILA after its oral supplementation to mice (n=4-11). Data represent mean ± SEM. *$p$ < 0.05, **$p$ < 0.01, ***$p$ < 0.001. two-tailed unpaired Student's $t$-test.

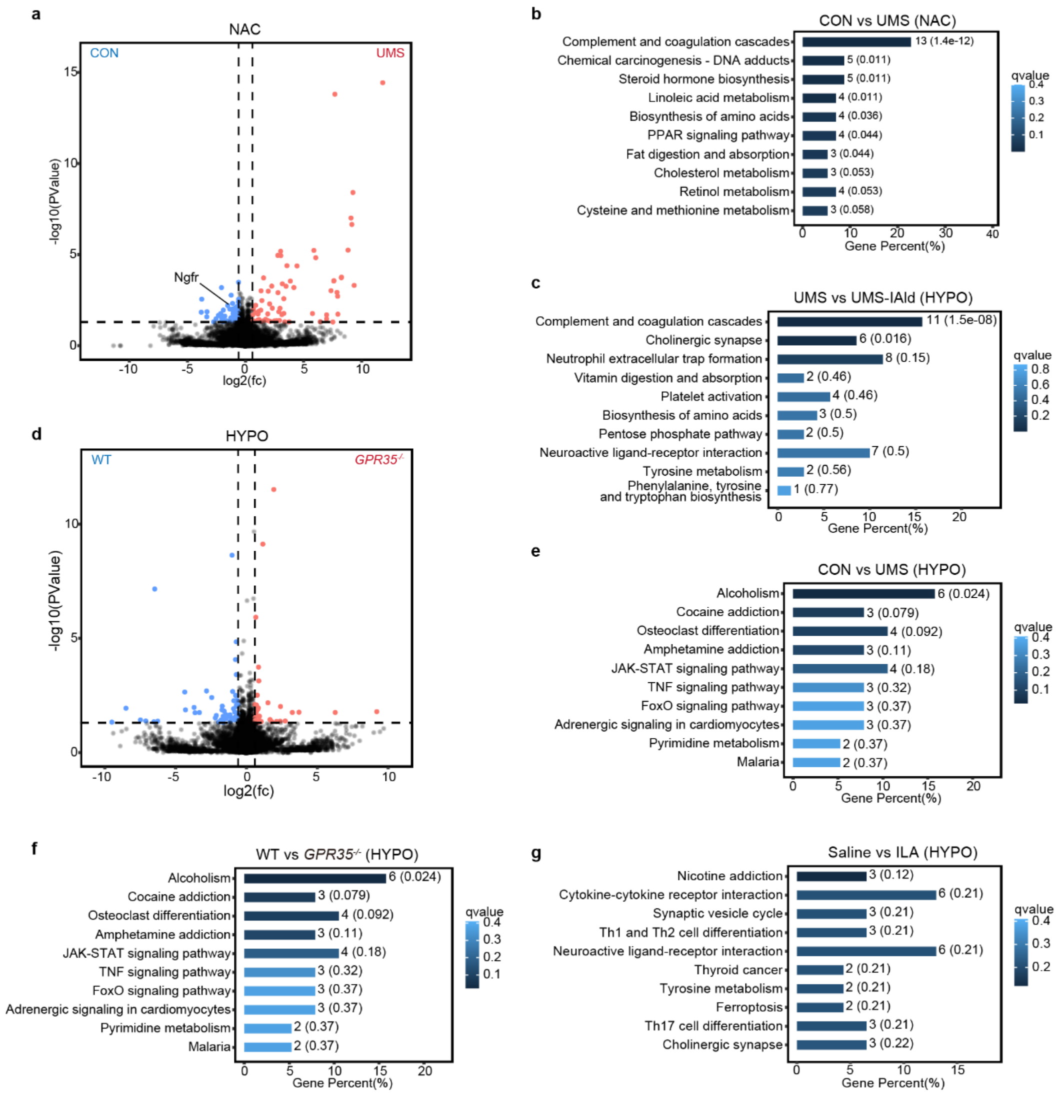


**Fig.S9 ILA and IAld counteractively regulate neuroplasticity.**

(a) Volcano plot displaying 122 differentially expressed genes (DEGs, log2 fold change >0.2, adjusted $p$ < 0.05) in the NAc from Con *vs* UMS mice (n=3 mice/group). (b) KEGG pathway analysis of differential biological processes in the NAc from Con *vs* UMS mice. (c)Top 10 significantly enriched KEGG pathways in the hypothalamus of UMS vs UMS+IAld group (n=3). (d) Volcano plot displaying 90 DEGs in the hypothalamus from WT vs *Gpr35*$^{-/-}$ mice. (e) Top 10 significantly enriched KEGG pathways in the hypothalamus of WT vs UMS group (n= 3 mice/group). (f) KEGG pathway analysis of differential biological processes in the hypothalamus from WT vs *Gpr35*$^{-/-}$ mice (n=3). (g) KEGG pathway analysis of differential biological processes in the hypothalamus from vehicle *vs* ILA-treated mice (n=3).

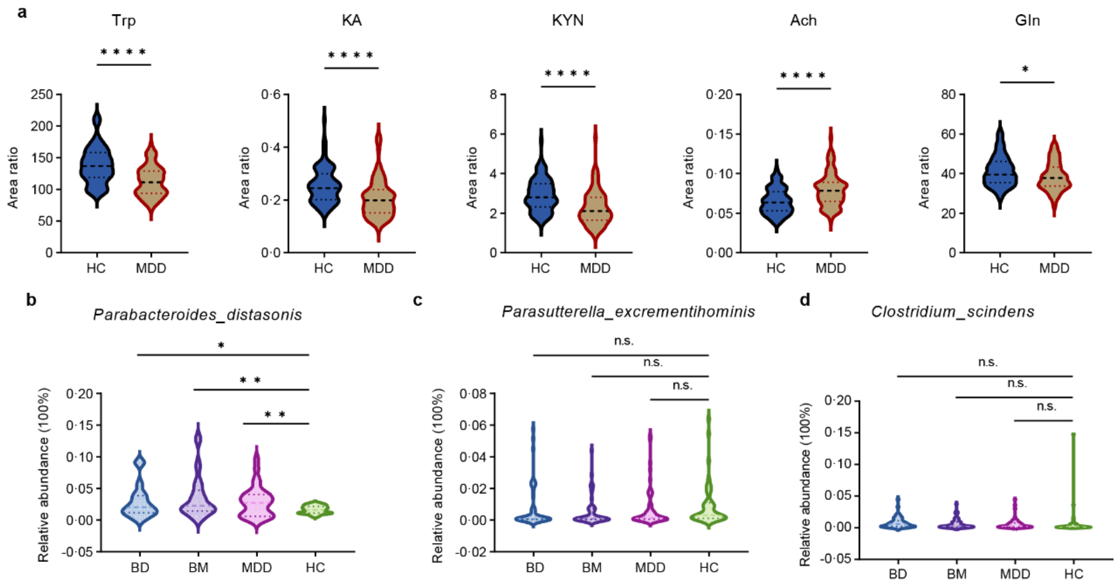


**Fig.S10 IAld is reduced in depressive patients.**

(a) Violin plot of the abundance of host tryptophan metabolites and neurotransmitters in the serum of healthy control (HC) and MDD patients. Trp, tryptophan; KA, kynurenic acid; KYN, kynurenine; Ach, acetylcholine; Gln, glutamine. (b) Relative abundance of *P.distasonis* in the gut microbiome of HC, MDD, bipolar disorder (BD) and bipolar mania (BM) patients assessed by metagenomic sequencing. (c,d) Relative abundance of *P.excrementihominis* and *C. scindens* in the gut microbiome of HC and MDD patients assessed by *16S* rRNA sequencing. *$p$ < 0.05, **$p$ < 0.01, ****$p$< 0.0001. n.s., no significance.

### Supplementary Tables

**Table S1 Summary of the stress schedule of the unpredictable mild stress procedure.**

| Day1 | Day2 | Day3 | Day4 | Day5 | Day6 | Day7 |
|---|---|---|---|---|---|---|
| Morning: confinement in tube for 30 min. Afternoon: confinement in tube for 30 min. Evening: crowding stress overnight. | Morning: cage shaking for 30 minutes. Afternoon: Tail suspension for 6 min; Noise stress for 30 min. Evening: damp bedding overnight. | Morning: attacked by CD-1 mice for 60 min. Evening: persistent illumination overnight. | Morning: Solitary cage feeding. Afternoon: tilted cage overnight. | Morning: tilted cage. Afternoon: bedding material deprivation overnight. | Morning: tilted cage and bedding material deprivation. Afternoon: Wind stress for 30 min (Blow dryer at high speed for 3min, rest for 3min as a cycle) | Morning: Each mouse was acclimated to the shock box for 1min, and the mice were randomly shocked 20 times for 5 s each over the next 6 min. |

**Table S2 Primer sequences for the qPCR analysis of mouse brain samples.**

| Gene | Forward(5'-3') | Reverse(5'-3') |
|---|---|---|
| *Gap43* | ATAACTCCCCGTCCTCCAAGG | GTTTGGCTTCGTCTACAGCGT |
| *Gpr35* | ACAACCTGTAACAGCACCCTC | GCGATAGCAGAATACCCAGAGT |
| *Tbr1* | GCAGCAGCTACCCACATTC | GTCCTTGGAGTCAGGAAAATTGT |
| *Gapdh* | CTCTCTGCTCCTCCTGTTCGAC | TGAGCGATGTGGCTCGGCT |
| *Glua1* | TCCCCAACAATATCCAGATAGGG | AAGCCGCATGTTCCTGTGATT |
| *Glua2* | TTCTCCTGTTTTATGGGGACTGA | CTACCCGAAATGCACTGTATTCT |
| *CamkIIα* | ACCTGCACCCGATTCACAG | TGGCAGCATACTCCTGACCA |
| *CamkIIβ* | TCACCGACGAGTACCAGCTA | GGCAGATCCGAGCTTCTCTC |
| *Bdnf* | TCATACTTCGGTTGCATGAAGG | AGACCTCTCGAACCTGCCC |
| *Gdnf* | AGAGGGGCAAAAATCGGGG | CCGCTGCAATATCGAAAGATCA |
| *Ngf* | CCAGTGAAATTAGGCTCCCTG | CCTTGGCAAAACCTTTATTGGG |
| *Grm4* | GACCGCATCAACAACGACC | GTGCCGTCCTTCTCGATGAG |

**Table S3 Primer sequences for the qPCR analysis of mouse intestinal samples.**

| Gene | Forward(5'-3') | Reverse(5'-3') |
|---|---|---|
| *Tjp1* | GCTTTAGCGAACAGAAGGAGC | TTCATTTTTCCGAGACTTCACCA |
| *Ocln* | TTGAAAGTCCACCTCCTTACAGA | CCGGATAAAAAGAGTACGCTGG |
| *Jam1* | TCTCTTCACGTCTATGATCCTGG | TTTGATGGACTCGTTCTCGGG |
| *Gapdh* | CTCTCTGCTCCTCCTGTTCGAC | TGAGCGATGTGGCTCGGCT |
| *Cldn1* | GCCTTGATGGTAATTGGCATCC | GGCCACTAATGTCGCCAGAC |
| *Cldn2* | CAACTGGTGGGCTACATCCTA | CCCTTGGAAAAGCCAACCG |
| *Cldn3* | ACCAACTGCGTACAAGACGAG | CGGGCACCAACGGGTTATAG |
| *Il6* | TAGTCCTTCCTACCCCAATTTCC | TTGGTCCTTAGCCACTCCTTC |
| *Il17* | TTTAACTCCCTTGGCGCAAAA | CTTTCCCTCCGCATTGACAC |
| *Il1β* | GAAATGCCACCTTTTGACAGTG | TGGATGCTCTCATCAGGACAG |
| *Tnfα* | ATCCTCTCTATCCTGCGACAC | GGGCTCTGGTTCTCAAACACT |

**Table S4. Primer sequences for qPCR analysis of bacterial strains.**

| Gene | Forward (5’-3’) | Reverse (5’-3’) |
|---|---|---|
| *P.distasonis* | GTACACACCGCCCGT | GTATGACCTCGGTACGGA |
| *P.excrementihominis* | AAGTAAAATTCTCAGTAACGCAGC | GCTCTCATTACAAGAGCTTCC |
| *C. scindens* | *AACTTTCATGGCGGACACAC* | *AATATCGCAGAGTTCCGGGT* |
| Bacterial *16S* rDNA | CCTACGGGAGGCAGCAG | ATTACCGCGGCTGCTGG |

**Table S5. Mass spectrometry detection parameters for tryptophan metabolites and neurotransmitters.**

| Metabolite | Detection mode | Retention time(min) | Q1(m/z) | Q3(m/z) |
|---|---|---|---|---|
| Trp | positive | 3.58 | 205.2 | 188.1 |
| Kyn | positive | 2.7 | 209.2 | 192.1 |
| KA | positive | 4.66 | 190.1 | 144.1 |
| 3-HK | positive | 3.98 | 225.2 | 208.1 |
| 5-HT | positive | 2.34 | 177.6 | 161.1 |
| 5-HIAA | positive | 6.58 | 192 | 146.2 |
| Gln | positive | 1.57 | 147.4 | 130.1 |
| Glu | positive | 1.43 | 148.2 | 84 |
| GABA | positive | 1.38 | 104.1 | 87 |
| NE | positive | 1.52 | 170 | 152 |
| Ach | positive | 1.33 | 146 | 87.4 |
| ILA | positive | 6.01 | 206.2 | 118 |
| IAM | positive | 5.24 | 175.2 | 118.1 |
| IAA | positive | 5.91 | 176.2 | 115.1 |
| IAld | positive | 5.76 | 146.2 | 130.1 |
| 1-MT | positive | 4.48 | 219 | 160 |

**Table S6. Mass spectrometry detection parameters for bile acid pathway metabolites.**

| Metabolite | Detection mode | Retention time(min) | Q1(m/z) | Q3(m/z) |
|---|---|---|---|---|
| dhCA | negative | 4.09 | 401.2 | 401.3 |
| LCA | negative | 27.57 | 375.2 | 375.2 |
| UDCA | negative | 13.94 | 391.283 | 391.283 |
| HDCA | negative | 15.17 | 391.284 | 391.284 |
| CDCA | negative | 25.26 | 391.285 | 391.285 |
| DCA | negative | 25.41 | 391.286 | 391.286 |
| CA | negative | 13.97 | 407.301 | 407.301 |
| αMCA | negative | 13.96 | 407.302 | 407.302 |
| βMCA | negative | 13.94 | 407.303 | 407.303 |
| GUDCA | negative | 24.91 | 448.301 | 448.301 |
| GCDCA | negative | 24.91 | 448.303 | 448.303 |
| GDCA | negative | 24.91 | 448.302 | 448.302 |
| GCA | negative | 14.43 | 464.301 | 464.301 |
| TLCA | negative | 26.34 | 481.8 | 481.8 |
| TUDCA | negative | 25.23 | 498.001 | 498.001 |
| THDCA | negative | 25.23 | 498.002 | 498.002 |
| TCDCA | negative | 25.23 | 498.003 | 498.003 |
| TDCA | negative | 25.23 | 498.004 | 498.004 |
| TCA | negative | 15.75 | 514.001 | 514.001 |
| TαMCA | negative | 9.18 | 514.002 | 514.002 |
| TβMCA | negative | 9.18 | 514.003 | 514.003 |
